# Tumor-responsive, multifunctional CAR-NK cells cooperate with impaired autophagy to infiltrate and target glioblastoma

**DOI:** 10.1101/2020.10.07.330043

**Authors:** Jiao Wang, Sandra Toregrosa-Allen, Bennett D. Elzey, Sagar Utturkar, Nadia Atallah Lanman, Victor Bernal-Crespo, Matthew M. Behymer, Gregory T. Knipp, Yeonhee Yun, Michael C. Veronesi, Anthony L. Sinn, Karen E. Pollok, Randy R. Brutkiewicz, Kathryn S. Nevel, Sandro Matosevic

**Affiliations:** Department of Industrial and Physical Pharmacy, Purdue University, West Lafayette, IN, USA; Center for Cancer Research, Purdue University, West Lafayette, IN, USA; Department of Comparative Pathobiology, Purdue University, West Lafayette, IN, USA; Histology Research Laboratory, Center for Comparative Translational Research, College of Veterinary Medicine, Purdue University, West Lafayette, IN, USA; Department of Radiology and Imaging Sciences, Indiana University School of Medicine, Indianapolis, IN, USA; In Vivo Therapeutics Core, Indiana University Melvin and Bren Simon Comprehensive Cancer Center, Indiana University School of Medicine, Indianapolis, IN, USA; Department of Pharmacology and Toxicology, Indiana University School of Medicine, Indianapolis, IN, USA; Department of Pediatrics, Herman B Wells Center for Pediatric Research, Indiana University School of Medicine, Indianapolis, IN, USA; Department of Medical and Molecular Genetics, Indiana University School of Medicine, Indianapolis, IN, USA; Department of Microbiology and Immunology, Indiana University School of Medicine, Indianapolis, IN, USA; Department of Neurology, Indiana University School of Medicine, Indianapolis, Indiana, USA

**Keywords:** glioblastoma, NK cells, autophagy, CD73, chimeric antigen receptor, cancer immunotherapy, immunometabolism, tumor microenvironment

## Abstract

Tumor antigen heterogeneity, a severely immunosuppressive tumor microenvironment (TME) and lymphopenia resulting in inadequate immune intratumoral trafficking have rendered glioblastoma (GBM) highly resistant to therapy. As a result, GBM immunotherapies have failed to demonstrate sustained clinical improvements in patient overall survival (OS). To overcome these obstacles, here we describe a novel, sophisticated combinatorial platform for GBM: the first multifunctional immunotherapy based on genetically-engineered, human NK cells bearing multiple anti-tumor functions, including local tumor responsiveness, that addresses key drivers of GBM resistance to therapy: antigen escape, poor immune cell homing, and immunometabolic reprogramming of immune responses. We engineered dual-specific CAR-NK cells to bear a third functional moiety that is activated in the GBM TME and addresses immunometabolic suppression of NK cell function: a tumor-specific, locally-released antibody fragment which can inhibit the activity of CD73 independently of CAR signaling and decrease the local concentration of adenosine. The multifunctional human NK cells targeted patient-derived GBM xenografts, demonstrated local tumor site specific activity in the tissue and potently suppressed adenosine production. We also unveil a complex reorganization of the immunological profile of GBM induced by inhibiting autophagy. Pharmacologic impairment of the autophagic process not only sensitized GBM to antigenic targeting by NK cells, but promoted a chemotactic profile favorable to NK infiltration. Taken together, our study demonstrates a promising new NK cell-based combinatorial strategy that can target multiple clinically-recognized mechanisms of GBM progression simultaneously.

## Introduction

Glioblastoma (GBM) is the most common and deadliest malignant type of primary brain tumor in adults and children^1,2^. GBM patients are poorly responsive to traditional treatments, resulting in a grim prognosis that has only modestly improved over the past several decades, motivating the hunt for new treatment approaches^3,4^. So far, chimeric antigen receptor-engineered NK (CAR-NK) cells targeting single GBM antigens―EGFR, EGFRvIII or ErbB2/HER2―have been limited to the use of NK cell lines, and the overall response rates have been disappointingly low and inconsistent^5–7^. These responses appear to mirror the clinical hurdles of single antigen targeted CAR-T therapies for GBM^8–12^. CAR-T cells, administered to target single GBM antigens *via* intracavitary, intraventricular or intravenous routes, have so far resulted in inconclusive durable responses^10^.

Pre-clinical and patient data have pointed to the heterogeneity of the GBM tumor microenvironment (TME) as a uniquely complex obstacle to overcome^13–15^. This is reflected in immunotherapies tested so far having failed to improve GBM patient overall survival (OS) in phase III clinical trials^16–18^. GBM induces localized lymphopenia to drive disease progression and resist treatment. In addition, the tumor’s heterogeneity is broad, with each of the known GBM subtypes—classical, mesenchymal, neural and proneural—displaying diverse genetic and epigenetic signatures associated with distinct and variable cell plasticities^19,20^. Not surprisingly, outgrowth of antigen escape variants has been recorded clinically with most GBM-associated antigens to date^9,23,24^, resulting in immune evasion and resistance to treatment^25^. And though strategies including dual antigen-targeting or programmable, tumor-sensing CARs—so far primarily in the context of adoptive T cell therapy— have been evaluated pre-clinically to combat such evasion, GBM employs mechanisms beyond antigen escape to avoid targeting. Treatment evasion by GBM is fueled by a heavily immunosuppressive, hypoxic TME which provides a niche unfavorable to NK cell effector function^26,27^. A subset of GBM cells, glioma stem-like cells (GSCs), contribute to treatment resistance and are poorly recapitulated by conventional GBM model cell lines, including U87MG^28^. Metabolic and functional pathways, moreover, converge to fuel the tumor’s invasiveness by driving, for instance, exhaustion of immune cells^29,30^. For instance, immunometabolic dysregulation of NK cell function in GBM is driven in part by the activity of ecto-5′-nucleotidase (CD73). CD73 is a hypoxic ectoenzyme that we and others have found to be associated with a negative prognosis^31,32^, and has emerged as an attractive clinical target^33^. Moreover, we have previously shown that the CD73-driven accumulation of extracellular adenosine (ADO)^34,35^ leads to significant purinergic signaling-mediated impairment of NK cell activity^36^.

Although they are among the most abundant lymphocytes found within the GBM TME, NK cells are still present in insufficient amounts in these tumors and exhibit a highly dysfunctional phenotype^37–39^. The need to recapitulate NK cell function lost to multiple complex mechanisms not only presents a significant challenge to traditional CAR-NK therapy, but requires a greater presence specifically within the tumor tissue to mount meaningful clinical responses. Our challenge has been to identify specific targets and processes that rescue NK dysfunction, enhance GBM tumor targeting, and increase NK cell recruitment into the TME.

Here, we describe the first multifunctional, engineered human NK cell-based therapy for glioblastoma developed around the programmed targeting of three clinically relevant pathways of GBM progression: antigen escape, immunometabolic suppression and poor intratumoral NK cell presence. First, we achieved dual antigen recognition by modifying NK cells with multi-CARs to target disialoganglioside (GD2) and ligands to natural killer group 2D (NKG2D), which are widely expressed on human GBM^40–42^. Within the same NK cells expressing the GD2- and NKG2D-based CARs, we then engineered the concomitant local release of an antibody fragment that impairs immunosuppressive purinergic signaling by blocking the activity of CD73 *via* the cleavage of a tumor-specific linker. The release of the linker is dependent on the activity of proteases that are upregulated in the tumor microenvironment^43^. Such local release is able to avoid systemic toxicities owing to its tumor-specific activation that occurs independently of CAR-based signaling. This is the first example of fully-engineered, multifunctional NK cells that simultaneously target immunometabolic reprogramming of immune responses and immune-evasion pervasive in the GBM TME.

We report the homing of such multifunctional NK cells was enhanced when administered in conjunction with autophagy inhibitors in patient-derived GBM xenografts. Autophagy inhibition triggered significant NK cell chemotaxis alongside upregulated CCL5 and CXCL10 gradients, while at the same time decreasing amounts of tumor-promoting CCL2 and CXCL12. Disabling autophagy further revealed sophisticated and complex reorganization of anti-GBM immunological responses which could contribute to enhanced CAR-NK effector function. Clinical efficacy of adding an autophagy inhibitor to GBM therapy was demonstrated in a prospective controlled randomized trial, where chronic administration of chloroquine (CQ), a common, FDA-approved autophagy inhibitor demonstrated a significantly-enhanced response of GBM to antineoplastic therapy. Treatment with CQ also sensitized cancer cells to exogenous agents, indicating that such treatments are clinically safe and well-tolerated^44,45^. We reveal, for the first time, a nuanced and potentially important role for autophagy inhibitors such as CQ in adoptive human CAR-NK therapy through the first direct examination of the effect of targeting autophagy on CAR-NK cell immunotherapy against GBM.

These studies aim to expand the repertoire of GBM-targeting NK cell-based immunotherapy, and describe a first example of addressing, simultaneously, the challenge of tumor antigen heterogeneity, an immunosuppressive TME and insufficient intratumoral trafficking of NK cells.

## Results

### NK cells engineered with a multifunctional, responsive CAR-based construct can target multiple antigens widely present in human GBM

We analyzed the correlation between the expression profiles of genes corresponding to NK cells and a series of functional genes encoding various pro- or anti-tumorigenic ligands and targets in GBM using an RNAseq patient dataset (TCGA) to infer the relationship between expression of these functional genes and NK cell presence. These include the NK cell signature gene set (*NCR1, NCR3, KLRB1, CD160* and *PRF1*), *NT5E* (which encodes CD73), *B4GALNT1* (which encodes the enzyme producing GD2), *MICA*/*B* (which encode common ligands for NKG2D), as well as *CCL5* and *CXCL10* (encoding two eponymous chemokines). *NT5E* and *B4GALNT1* had a negative correlation with individual genes representing the NK signature set in GBM (r < 0), suggesting that the antigens encoded by these genes act to suppress NK cell function and proliferation in GBM. Alternatively, genes which are known to drive NK effector responses, including NKG2D-ligand encoding *MICA*/*B*, as well as *CCL5* and *CXCL10* revealed a positive correlation with individual NK signature genes (r > 0) (**Figure 1A**). Classification of GBM patients using TCGA RNA-seq data^46^ into high/low groups based on 50% upper and lower quartiles identified 151/156 patients overexpressing at least one of four targeted genes (*NT5E*, *B4GALNT1*, *MICA* or *MICB*). Enrichment analysis against the NK signature gene set using GBM patients’ data with high expression of specific genes indicate a negative correlative expression (NES < 1) for genes *NT5E* and *B4GALNT1*, and a positive correlative expression (NES > 1) for genes *MICA/MICB*, *CXCL10* and *CCL5* (**Figure 1B**; **Table S1**). A Venn diagram showing the association of gene expression among patient numbers indicates a corresponding and expected lower number of patients expressing multiple gene combinations (**Figure 1C** and **D**).

**Figure 1.**
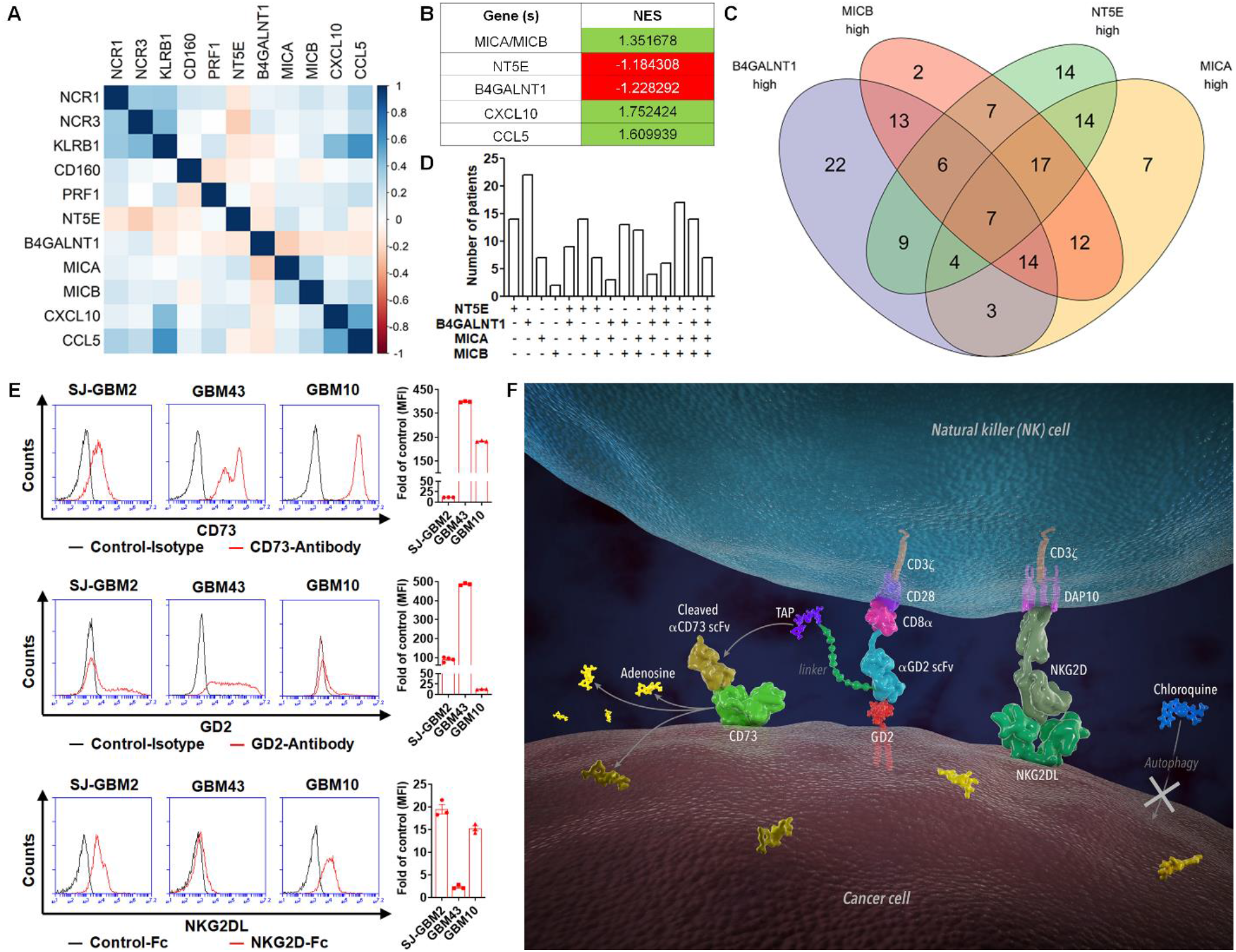
Correlative analysis of gene expression in GBM patient data, surface expression of CD73, GD2 and NKG2DL in patient-derived GBM, and the design of the multifunctional NK-based GBM immunotherapy. (**A**) Correlation between normalized expression (FPKM) of selected genes using data from 156 GBM patients. Pearson's correlation coefficients are shown with continuous gradient colors. (**B**) Correlation between normalized expression (FPKM) of the entire NK gene set and individual genes. Correlation expressed as normalized enrichment scores (NES). (**C**) Venn diagram showing the number of GBM patients with high expression of at least one of the given 4 genes, *NT5E, B4GALNT1*, *MICA* and *MICB*. (**D**) Bar graph showing the patient distribution of gene expression in GBM tumors based on the four genes in (**C**). (**E**) Surface expression of CD73, GD2 and NKG2DL on different types of patient-derived GBM cells including SJ-GBM2 (pediatric), GBM43 (primary) and GBM10 (recurrent) determined by flow cytometry. Herein, results are reported as fold-change (FC) over control. Data are shown as mean ± SEM. (**F**) A schematic illustration showing the multifunctional, tumor-responsive engineered NK cells and their working mechanisms. We developed an innovative approach to improve NK cell-based immunotherapy for GBM *via* 1) generation of engineered multifunctional NK cells which consist of a tumor-specific, locally-cleavable single chain antibody targeting CD73 alongside dual chimeric antigen receptors directed against GD2 and NKG2D ligands and 2) inhibition of autophagy in GBM cells to promote NK cell recruitment and GBM sensitization to CAR-NK therapy. Note: TAP stands for tumor associated protease. aCD73 scFv stands for anti-CD73 scFv. aGD2 scFv stands for anti-GD2 scFv.

The two pro-tumorigenic markers GD2 and CD73 are widely and heterogeneously expressed on different types of GBM, including patient-derived primary adult (GBM43), pediatric (SJ-GBM2), and recurrent adult (GBM10) brain tumor cells (**Figure 1E**). Among these, the ectoenzyme CD73 was previously demonstrated to be a negative prognostic factor^32^. Conversely, cognate ligands to the potent activating NK cell receptor NKG2D are also highly present on GBM^41^, though NKG2D is often downregulated in the TME (**Figure S1**)^47^, supporting the need for increasing its expression on NK cells through engineering.

To achieve responsive and combinatorial targeting of these GBM antigens, we first engineered a bicistronic vector to express two individual CARs simultaneously, yielding NK cells expressing both a GD2.CD28.CD3ζ CAR and NKG2D.DAP10.CD3ζ CAR. The GD2.CD28.CD3ζ CAR was then further engineered in tandem, following a cleavable, tumor-sensitive linker, with an anti-CD73 scFv to generate CD73-GD2.CD28.CD3ζ-CAR (**Figure S2**). This resulted in the localized release of a CD73-blocking antibody fragment in the GBM TME independent of CAR activation. Combining this multifunctional, responsive engineered NK cell approach is the adjuvant administration of an autophagy inhibitor (i.e. CQ) to both boost NK cell infiltration more deeply into the GBM TME and reorganize the immunological responses of GBM in favor of CAR-based targeting (**Figure 1F**).

Functionality of individual genetic components of the multifunctional construct was evaluated by generating separate versions of each element, including NKG2D.DAP10.CD3ζ-CAR (**Figure 2A**), GD2.CD28.CD3ζ-CAR (**Figure 2F**) and the multifunctional CAR bearing the full construct (**Figure 2K**). Each CAR was separately expressed into NK-92 cells and expression of the corresponding CAR structure was verified (**Figure 2B**, **G** and **L**). In all cases (**Figure 2C**, **H** and **M**), the CAR-expressing NK cells retained high viability (>95%). At the same time (**Figure 2D**), robust NKG2D.DAP10.CD3ζ-CAR expression on NK cells was achieved (~44%). NKG2D.DAP10.CD3ζ-CAR-NK92 cells showed higher cytotoxic activity of GBM43 cells compared to non-transfected cells (**Figure 2E**). Similar expression was achieved with (**Figure 2I**), the GD2.CD28.CD3ζ-CAR (~42%), with a correspondingly higher cytotoxicity against patient-derived GBM target cells (**Figure 2J**). This confirmed that the GD2.CD28.CD3ζ-CAR construct was capable of effectively recognizing and eliminating GD2-expressing GBM cells. When engineered to express the complete multifunctional sequence, NK-92 cells showed a significant increase in NKG2D expression (**Figure 2N**), anti-GD2 scFv expression and anti-CD73 scFv expression (**Figure 2O**). Compared to NK-92 cells, primary NK (pNK) cells derived from the peripheral blood of healthy donors generally possess higher lytic activity against patient-derived GBM target cells, as we observed. We isolated pNK cells from healthy adult donors (**Figure 2P** and **S3**) and engineered them by lentiviral transduction in the presence of DEAE-dextran to express the full multifunctional construct (CD73.mCAR-pNK). After two rounds of transduction, an expected, donor-specific drop in viability of the donor NK cells due to the transduction procedure (**Figure 2Q** and **S3**) was accompanied with significantly increased expression of NKG2D, anti-GD2 scFv and anti-CD73 scFv (**Figure 2R**, **S** and **S3**). Moreover, CD73.mCAR-pNK cells were able to expand substantially over 2-weeks in commercial NK-MACS^®^ medium without the use of feeders (**Figure 2T**) while retaining stable gene expression. Collectively, these results validate the use of CD73, GD2 and NKG2DL as combinatorial GBM targets, and provides a rationale for investigating their use for NK cell-based GBM immunotherapy.

**Figure 2.**
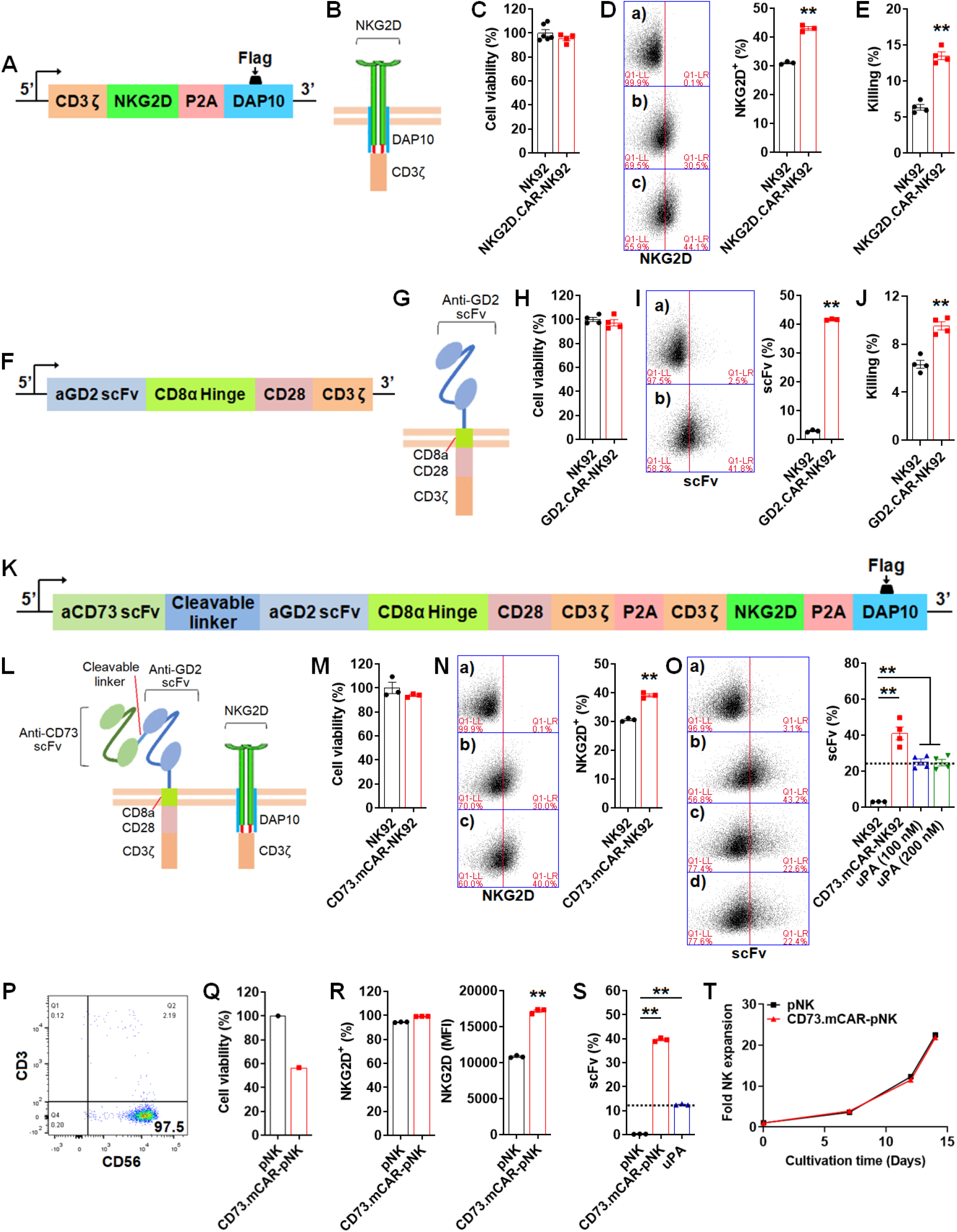
Generation of multifunctional genetically-engineered NK cells. (**A**) Schematic representation of transgene for NKG2D.DAP10.CD3ζ-CAR constructs targeting NKG2D ligands (NKG2DL). (**B**) Schematic representation of NKG2D.DAP10.CD3ζ-CAR structure. (**C**) Cell viability (%) of NK-92 cells engineered to express the NKG2D-CAR construct 48 hours after transfection. (**D**) NKG2D expression determined by flow cytometry on transfected CAR-NK92 cells. (**E**) *In vitro* cytotoxicity of NKG2D.DAP10.CD3ζ-CAR-NK92 cells and non-transfected NK-92 cells against GBM43 cells at an E/T ratio of 5 over 4 h. (**F**) Schematic representation of transgene structure representing the GD2.CD28.CD3ζ-CAR construct targeting GD2. (**G**) Schematic representation of GD2.CD28.CD3ζ-CAR structure. (**H**) The cell viability (%) of NK-92 cells engineered to express the GD2.CD28.CD3ζ-CAR construct after 48 h transfection. (**I**) Anti-GD2 scFv expression determined by flow cytometry on transfected CAR-NK92 cells. (**J**) *In vitro* cytotoxicity of GD2.CD28.CD3ζ-CAR-NK92 cells and non-engineered NK-92 controls against GBM43 cells at an E/T ratio of 5 over 4 h. (**K**) Schematic representation of transgene representing the complete multi-functional construct: tumor-responsive anti-CD73 scFv-secreting dual-specific CAR targeting NKG2DL and GD2. (**L**) Schematic representation of tumor-responsive anti-CD73 scFv secreting dual-specific CAR. (**M**) The cell viability (%) of NK-92 cells engineered to express the full multifunctional construct after 48 h transfection. (**N**) NKG2D expression determined by flow cytometry after 48 h transfection. (**O**) Expression of anti-CD73 scFv and anti-GD2 scFv on NK cells determined by flow cytometry following proteolytic cleavage with uPA to release the anti-CD73 ScFv fragment. (**P**) Flow cytometry data showing the purity of isolated peripheral blood-derived NK (pNK) cells (CD56^+^CD3^−^). (**Q**) The cell viability (%) of pNK cells engineered to express the full construct after two rounds of lentiviral transduction. (**R**) NKG2D expression on engineered pNK cells determined by flow cytometry after two rounds of lentiviral transduction. (**S**) Expression of anti-CD73 scFv and anti-GD2 scFv on pNK cells determined by flow cytometry after two rounds of lentiviral transduction. (**T**) Fold expansion of non-transduced pNK and CD73.mCAR-pNK cells in NK MACS^®^ medium. Note: the data shown through (**P**)-(**T**) is for isolated pNK cells from a representative donor. Data are shown as mean ± SEM. \**p* < 0.05, \*\**p*< 0.01.

### Multifunctional engineered NK cells can target patient-derived GBM cells while sparing normal cells

Compared to control NK-92 cells, CD73.mCAR-NK92 cells showed significantly improved killing of GBM43 cells after 4 h at various E/T ratios (**Figure 3A**). Furthermore, co-culture with GBM43 cells stimulated CD73.mCAR-NK92 cells to induce degranulation as measured by cell surface CD107a expression (**Figure 3B**) and up-regulate IFN-γ secretion (**Figure 3C**). After treating CD73.mCAR-NK92 cells with uPA (100 nM) to liberate the anti-CD73 scFv, the number of killed GBM43 cells decreased (**Figure 3D**). Additionally, the cleaved anti-CD73 scFv resulted in significantly decreased production of extracellular adenosine by GBM43 cells (**Figure S4** and **Figure 3E**), indicating that the enzymatic ability of CD73 on GBM was impaired and confirming that the anti-CD73 scFv is functional and specific for CD73, and can abrogate adenosine accumulation.

**Figure 3.**
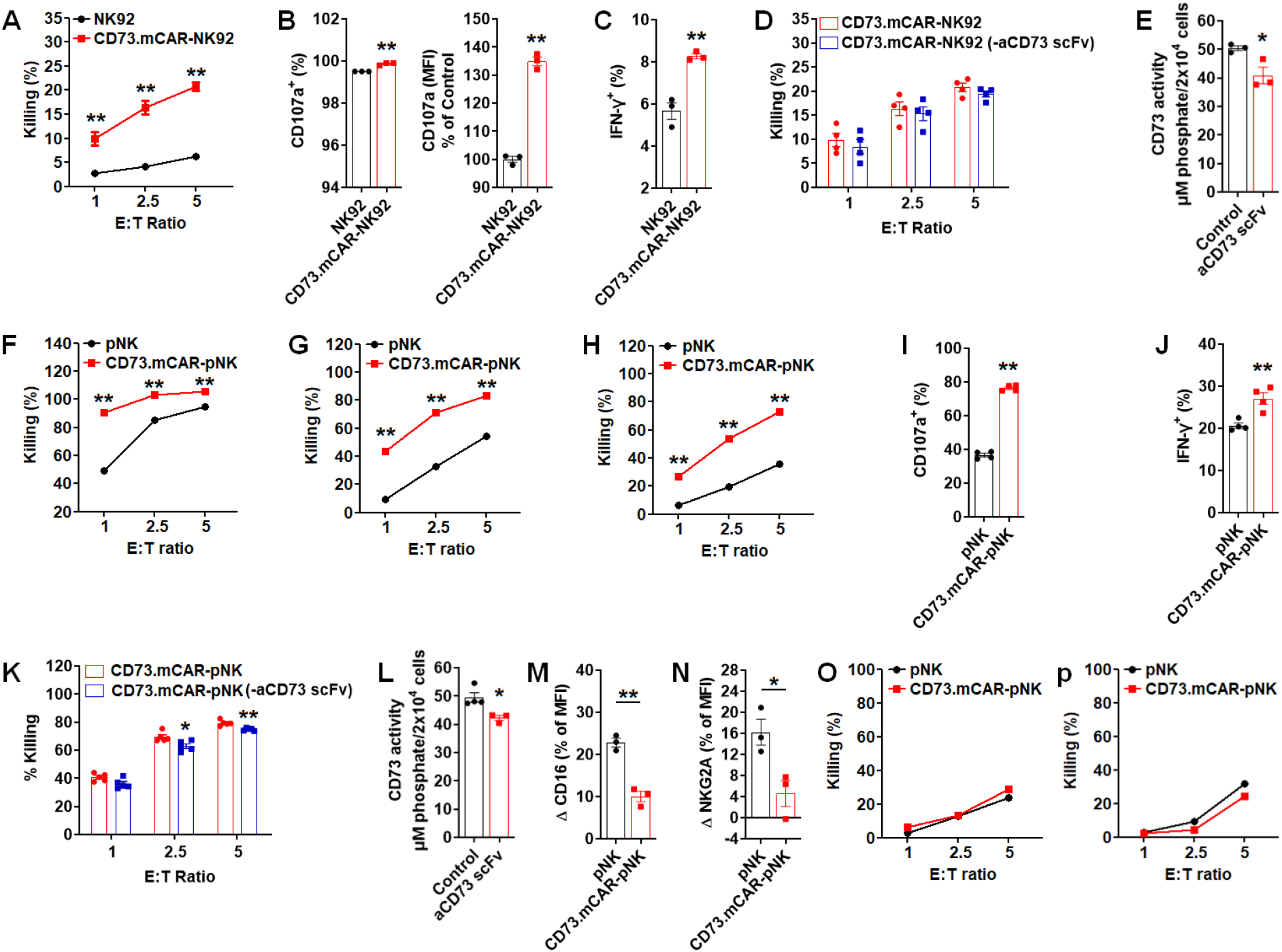
*In vitro* effector activity of multifunctional genetically-engineered NK cells against patient-derived GBM. (**A**) *In vitro* cytotoxicity of NK-92 and CD73.mCAR-NK92 cells against GBM43 cells at indicated E/T ratios over 4 h. (**B**) Degranulation of NK-92 and CD73.mCAR-NK92 cells [% CD107 and (MFI) CD107] after 4 h coculture with GBM43 cells (E/T ratio, 5:1). NK cells were analyzed by flow cytometry for surface CD107a expression as a marker of degranulation. (**C**) IFN-γ production by NK-92 and CD73.mCAR-NK92 cells (% IFN-γ) after 4 h coculture with GBM43 cells (E/T ratio, 5:1). (**D**) *In vitro* cytotoxicity of CD73.mCAR-NK92 and CD73.mCAR-NK92 (following aCD73 scFv cleavage) cells against GBM43 cells at indicated E/T ratios over 4 h. (**E**) CD73 activity of GBM43 cells after incubation with cleaved aCD73 scFv following release from uPA-treated CD73.mCAR-NK92 cells. (**F-H**) *In vitro* cytotoxicity of pNK and CD73.mCAR-pNK cells against different GBM cells, including SJ-GBM2, GBM43 and GBM10 cells, at indicated E/T ratios over 4 h. (**I**) Degranulation of pNK and CD73.mCAR-pNK cells (% CD107) after 4 h coculture with GBM43 cells (E/T ratio, 5:1). NK cells were analyzed by flow cytometry for surface CD107a expression as a marker of degranulation. (**J**) IFN-γ production of pNK and CD73.mCAR-pNK cells (% IFN-γ) after 4 h coculture with GBM43 cells (E/T ratio, 5:1). (**K**) *In vitro* cytotoxicity of pNK and CD73.mCAR-pNK (following aCD73 scFv cleavage) cells against GBM43 cells at indicated E/T ratios over 4 h. (**L**) CD73 activity of GBM43 cells after incubation with cleaved aCD73 scFv following cleavage from uPA-treated CD73.mCAR-NK cells. (**M**) Relative decrease in CD16 expression on pNK and CD73.mCAR-pNK cells (% of MFI) after 12 h coculture with GBM43 cells (E/T ratio, 5:1). (**N**) Relative increase in NKG2A expression on pNK and CD73.mCAR-pNK cells (% of MFI) after 12 h coculture with GBM43 cells (E/T ratio, 5:1). (**O** and **P**) *In vitro* cytotoxicity of pNK and CD73.mCAR-pNK cells against nonmalignant neural cell lines hCMEC/D3 and HCN-2 at indicated E/T ratios over 4 h. Note: the data shown through (**F**)-(**P**) is for isolated pNK cells in at least triplicates from a representative donor. Data are shown as mean ± SEM. \**p* < 0.05, \*\**p* < 0.01.

Human peripheral blood-derived NK cells, genetically-engineered to express the multi-functional construct (CD73.mCAR-pNK), resulted in efficient killing of SJ-GBM2 (pediatric), GBM43 (adult) and GBM10 (recurrent) cells, to significantly higher levels than those of control non-engineered NK cells (**Figure 3F-H**, **Figure S3**). Live imaging of the killing of GBM targets by native or engineered NK cells demonstrates the dynamic nature of this process and a higher killing specificity of GBM cells by CD73.mCAR-pNK cells (**Figure S5** and **Video 1**) compared to that by native human NK cells (**Figure S5** and **Video 2**). Stimulation by GBM cells contributed to a significantly increased NK cell degranulation and intracellular production of IFN-γ by CD73.mCAR-pNK cells (**Figure 3I** and **J**). CD73.mCAR-NK cells lacking the anti-CD73 scFv following uPA treatment displayed significantly decreased cytotoxic ability of target GBM43 cells after co-culture at effector/target (E/T) ratios of 2.5 and 5 for 4 h (**Figure 3K**). In addition, after treatment with anti-CD73 scFv cleaved from CD73.mCAR-pNK cells, GBM43 cells showed a significantly reduced ability to produce adenosine due to the loss of active CD73 (**Figure 3L**). The anti-tumor specificity imparted upon NK cells by genetic expression of the multifunctional construct resulted in enhanced resistance to the loss of CD16 upon contact with GBM cells (**Figure 3M**), as well as a reduced upregulation of NKG2A (**Figure 3N**) compared to changes observed on non-engineered human NK cells in response to GBM.

In order to address potential off-target effects due to multispecific targeting of any potential expression of these antigens on non-tumor tissues, we evaluated the ability of CD73.mCAR-pNK cells to target healthy cells. CD73.mCAR-pNK cells did not preferentially kill normal cells, specifically those belonging to neural lineages, including hCMEC/D3 and HCN-2 cells. CD73.mCAR-pNK cells exhibited effector activity comparable to that of control non-engineered pNK cells against healthy brain cells (**Figure 3O** and **P**). Although multiple mechanisms may be involved in the targeting of such cells⎯including a distinct pattern of inhibitory ligand expression, the low cytotoxicity rates induced by CD73.mCAR-pNK cells could be due to the relatively lower expression profiles for the targeted ligands on these cells compared with GBM cells (**Figure S6**). Together, these observations indicated the functional superiority of CD73.mCAR-pNK cells, and represent a promising path forward for highly specific anti-GBM immunotherapies without causing observable toxicities to normal cells.

### Functional targeting of autophagy in GBM sensitizes the tumor to enhanced NK activity and homing

Autophagy is considered a critical cell survival mechanism leading to and driving cancer development and progression^48^. In order to tamper with its ability to drive GBM resistance to therapy and establish its sensitization of GBM to immune cell treatment, we investigated the effects of blocking autophagy in GBM, either genetically, by targeting the *BECN1* gene through generating knockdown patient-derived GBM cells (*BECN1*^−^ GBM43), or pharmacologically, *via* treatment with chloroquine (CQ), a common, FDA-approved autophagy inhibitor (**Figure S7**). We tested these targeting approaches on NK cell infiltration and effector functions. CQ impaired viability and proliferation of GBM43 cells in a dose-dependent manner after 24 h (IC_50_ = 67.10 μM; **Figure 4A** and **B**).

**Figure 4.**
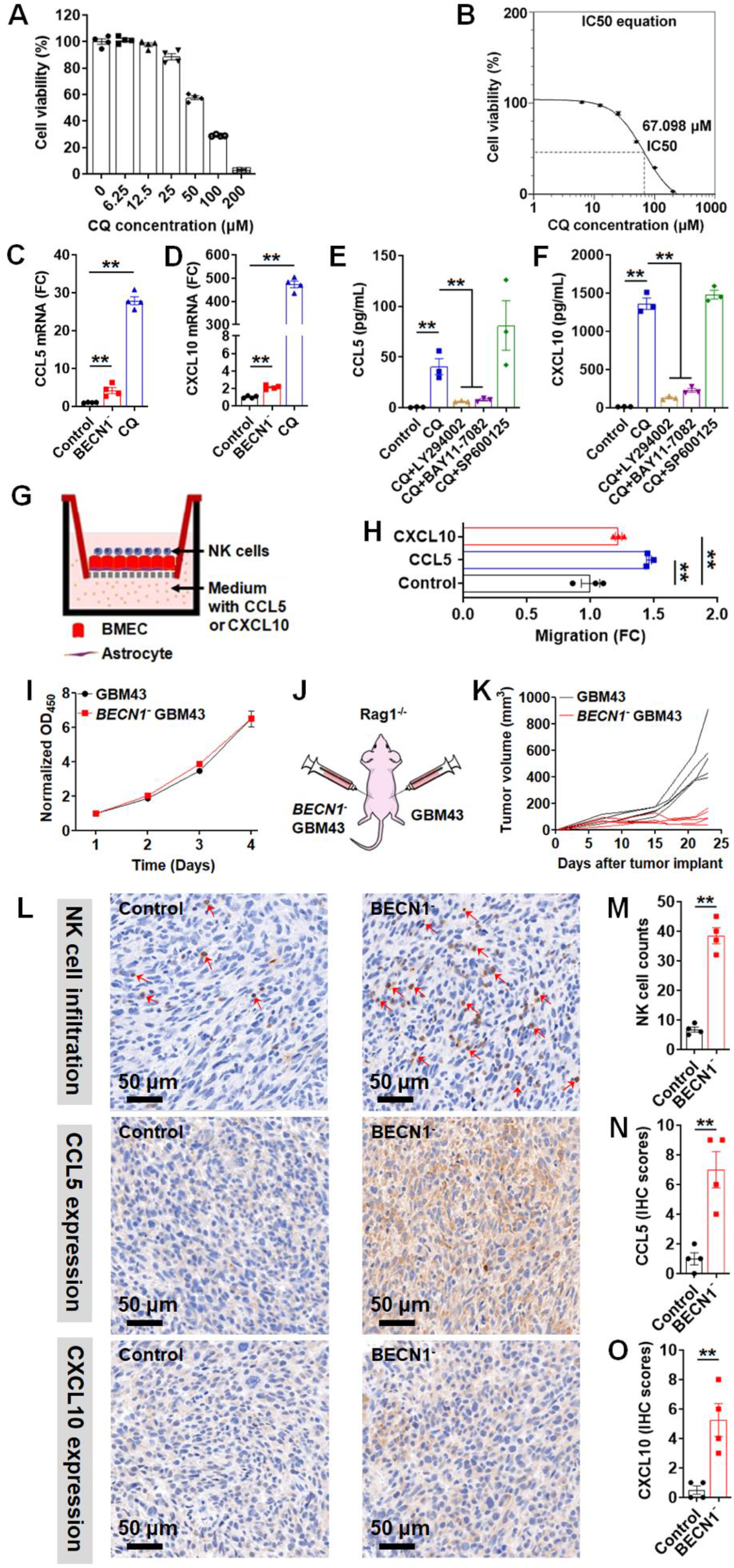
Effects of targeting autophagy in GBM on NK cell function and homing. (**A**) Viability of GBM43 cells after treatment with various concentrations of CQ for 24 h *in vitro*. (**B**) Cell viability (%) was plotted versus the Log [CQ concentration (μM)] and the IC_50_ was computed using Prism 5 (GraphPad Software Inc.) (**C** and **D)** Expression of CCL5 and CXCL10 mRNA in GBM43 cells transfected with *BECN1* shRNA lentivirus (*BECN1*^−^ GBM43) or in cells treated with CQ. Results are reported as a fold-change (FC) compared to non-transfected controls. (**E** and **F**) Quantification of CCL5 and CXCL10 in the supernatant of GBM43 cells following treatment with CQ for 24 h in the presence or absence of various inhibitors, including LY294002, BAY11-7082 and SP600125 by ELISA. (**G**) Schematic of *in vitro* experimental bilayer BBB model representing the experimental setup used to evaluate the effects of CCL5 or CXCL10 on pNK cell migration across the BBB. (**H**) Transmigration of pNK cells through experimental BBB. (**I**) *In vitro* viability of GBM43 and *BECN1*^−^ GBM43 cells as determined by CCK-8 assay. (**J**) Schematic of the experimental design to evaluate the effects of targeting autophagy through *BECN1* knockdown on tumor growth and NK cell infiltration. GBM43 cells were injected into right flank and *BECN1*^−^ GBM43 cells were injected into the left flank of Rag1^*−/−*^ mice. (**K**) Tumor growth was monitored and recorded on the indicated days. (**L**) Immunohistochemical (IHC) staining of NK cells (top row), CCL5 (middle row) or CXCL10 (bottom row) performed on indicated tumor sections using anti-NKp46, anti-CCL5 and anti-CXCL10 antibodies, respectively. Bar = 50 μm; 200× magnification. (**M**) Quantification of NK cells infiltrating control and *BECN1*^*−*^ tumors. Here, cell counts were recorded in 4 consecutive high-power fields (HPFs) at 200× magnification. (**N** and **O**) IHC scores of CCL5 and CXCL10 in control and *BECN1*^*−*^ tumors. Note: the data shown in (**H**) is for isolated pNK cells from a representative donor. Data are shown as mean ± SEM. \**p* < 0.05, \*\**p*< 0.01.

The gradient between chemokines and their cognate receptors initiates the directional movement of cells to sites with higher concentrations to ultimately regulate immune cell trafficking to tumors^49,50^. Due to the paucity of insight into chemokine-mediated trafficking of NK cells to GBM, we explored the consequence of inhibiting autophagy in GBM on the expression levels of a number of chemokines known to be associated with NK cell trafficking to tumors. Both *BECN1*^−^ GBM43 and CQ-treated GBM43 cells resulted in a substantial upregulation of CCL5 and CXCL10 at the transcriptional level as measured by a significant increase in their mRNA (**Figure 4C** and **D**). At the same time, the chemokines CCL2 and CXCL12 showed decreased transcriptional levels on GBM43 cells upon inhibition of autophagy (**Figure S8**).

To evaluate changes in the protein levels of these chemokines and gain more insight into the underlying molecular mechanisms, we quantified their concentrations in the conditioned media of CQ-treated GBM43 cells, by ELISA, in absence and presence of various pharmacological inhibitors driving signaling pathways for these chemokines. This included the PI3K inhibitor LY294002, NF-κB inhibitor BAY11-7082 and JNK inhibitor SP600125. CQ-treated GBM43 cells secreted significantly higher amounts of both CCL5 and CXCL10 compared with untreated control cells (**Figure 4E** and **F**). The elevated production of these chemokines was significantly inhibited by LY294002 or BAY11-7082, but not by SP600125. This suggests a mechanistic link between enhanced expression of CCL5 and CXCL10 upon inhibition of autophagy on GBM and activation of the PI3K/NF-κB pathway. To determine involvement of CCL5 and CXCL10 in the migration of NK cells, we tracked the migration of activated pNK cells toward CCL5 and CXCL10 gradients using an *in vitro* BBB model setup by a direct contact co-culture of primary human astrocytes and hCMEC/D3 human cerebral microvessel endothelial cells (**Figure 4G**)^51^. As shown in **Figure 4H**, activated pNK cells had a significantly increased ability to migrate along CCL5 (100 ng/mL) as well as CXCL10 (100 ng/mL) ligand gradients. The viability and proliferation of *BECN1*^−^ GBM43 cells were unaffected by *BECN1* knockdown (**Figure 4I**).

The impact of *BECN1* knockdown on GBM tumor growth and NK cell infiltration was further evaluated *in vivo* in Rag1^*−/−*^ mice, which bear no mature B and T lymphocytes but possess a mature NK cell compartment (**Figure 4J**). Tumors from the left flanks of mice engrafted with *BECN1*^−^GBM43 cells exhibited significantly slower growth and a smaller size than those arising from control GBM43 cells (**Figure 4K**). Immunohistochemical (IHC) staining was performed on extracted tumors to detect the expression of NKp46 as a measure of murine NK cell presence, alongside staining for CCL5 and CXCL10. A significantly higher expression of NKp46 was detected in tumors lacking *BECN1*, demonstrating a deeper infiltration of NK cells into GBM lacking the ability to perform autophagy compared with control tumors (**Figure 4L** and **M**). In addition, NK cells in *BECN1*^−^ tumors showed a higher distribution both at the tumor periphery and in intratumoral areas (**Figure S9**). The expression of CCL5 and CXCL10 was upregulated from relatively low baseline levels in control tumors to significantly higher levels in *BECN1*^−^ tumors (**Figure 4L**, **N** and **O**).

In addition to the increase in NK cell intratumoral infiltration, we found that CQ-mediated inhibition of autophagy had an obvious and significant effect on the expression of NKG2DL, CD73 and GD2 on GBM43 cells. Specifically, NKG2DL expression (both % and MFI) on GBM43 cells increased significantly after treatment with CQ at various concentrations for 24 h (**Figure S10A**). On the other hand, expression of CD73 (MFI) decreased following CQ treatment, even at a low concentration of 6.25 μM (~70% of control) (**Figure S10B**). In addition, production of adenosine in human GBM cells mediated by CD73 was also significantly reduced after treatment with CQ (**Figure S10C**). This effectively decreased the concentration of CD73 that needs to be deactivated by the NK cells. Expression of GD2 (MFI) decreased after treatment with CQ at concentrations of 50 μM or higher one. Conversely, the percentage expression (%) of GD2 decreased in response to up to 100 μM CQ, after which it remained constant (~46%) compared to that of control cells (~58%) (**Figure S10D**). To further investigate the impact of the inhibition of autophagy on the cytotoxic capacity of NK cells, we incubated CQ-treated GBM43 cells or *BECN1*^−^ GBM43 cells with pNK cells for 4 h at an E/T ratio of 5. We found that treatment with CQ sensitized GBM cells to superior killing by pNK cells (**Figure S10E** and **F**).

Together, our data demonstrated that targeting autophagy improves the infiltration of NK cells into the GBM tumor bed aided by higher levels of the chemokines CCL5 and CXCL10. In addition, disabling autophagy can also potentiate NK cell-mediated cytotoxicity against GBM cells by reprogramming their phenotypic signatures in favor of enhanced NK cell functionality, such as increased NKG2DL expression. The complex and unique reprogramming of the GBM TME induced by targeting autophagy can further sensitize treatment of this tumor with NK cells engineered to exploit these reprogrammed pathways.

### Multifunctional engineered NK cells efficiently target patient-derived GBM tumors *in vivo*

To evaluate the *in vivo* antitumor activity of multifunctional engineered NK cells as well as the effects induced by targeting autophagy on GBM, we first established a subcutaneous xenograft model by engrafting patient-derived GBM43 cells into NSG mice. The treatment schedule is summarized in **Figure 5A**. Briefly, 3 × 10^6^ GBM43 cells were subcutaneously (SC) implanted into the right flank of the mice (Day 0). 10 days later (Day 10), mice in the CQ group were intraperitoneally (IP) injected with CQ at 50 mg/kg for 3 weeks, once a week. One day later (Day11), the mice in pNK and CD73.mCAR-pNK groups were treated with 5 × 10^6^ adoptively-transferred pNK or CD73.mCAR-pNK cells intravenously (IV), once a week for 3 weeks. Starting on the day of the first injection of NK cells, all mice received 0.5 μg of IL-15 once every 3-4 days IP. Treatment with pNK cells significantly delayed tumor growth compared with mice in the PBS control group (mean tumor volume: 815.75 mm^3^ vs 1475 mm^3^, \*\**p*< 0.01; mean tumor weight: 1.13 g vs 2.31 g, \*\**p*< 0.01; **Figure 5B** and **C**). Compared to pNK cell-treated mice, CD73.mCAR-pNK cells mediated more potent antitumor responses and significantly reduced the tumor growth rate (mean tumor volume: 401.5 mm^3^ vs 815.75 mm^3^, \*\**p*< 0.01; mean tumor weight: 0.59 g vs 1.13 g, \*\**p*< 0.01). CQ treatment also showed significant inhibition of tumor growth compared with the PBS group (mean tumor volume: 953.25 mm^3^ vs 1475 mm^3^, \**p*< 0.05; mean tumor weight: 1.16 g vs 2.31 g, \*\**p*< 0.01; **Figure 5B** and **C**). There was no significant decrease in body weight of the mice in all groups throughout the entire treatment period (**Figure 5D**).

**Figure 5.**
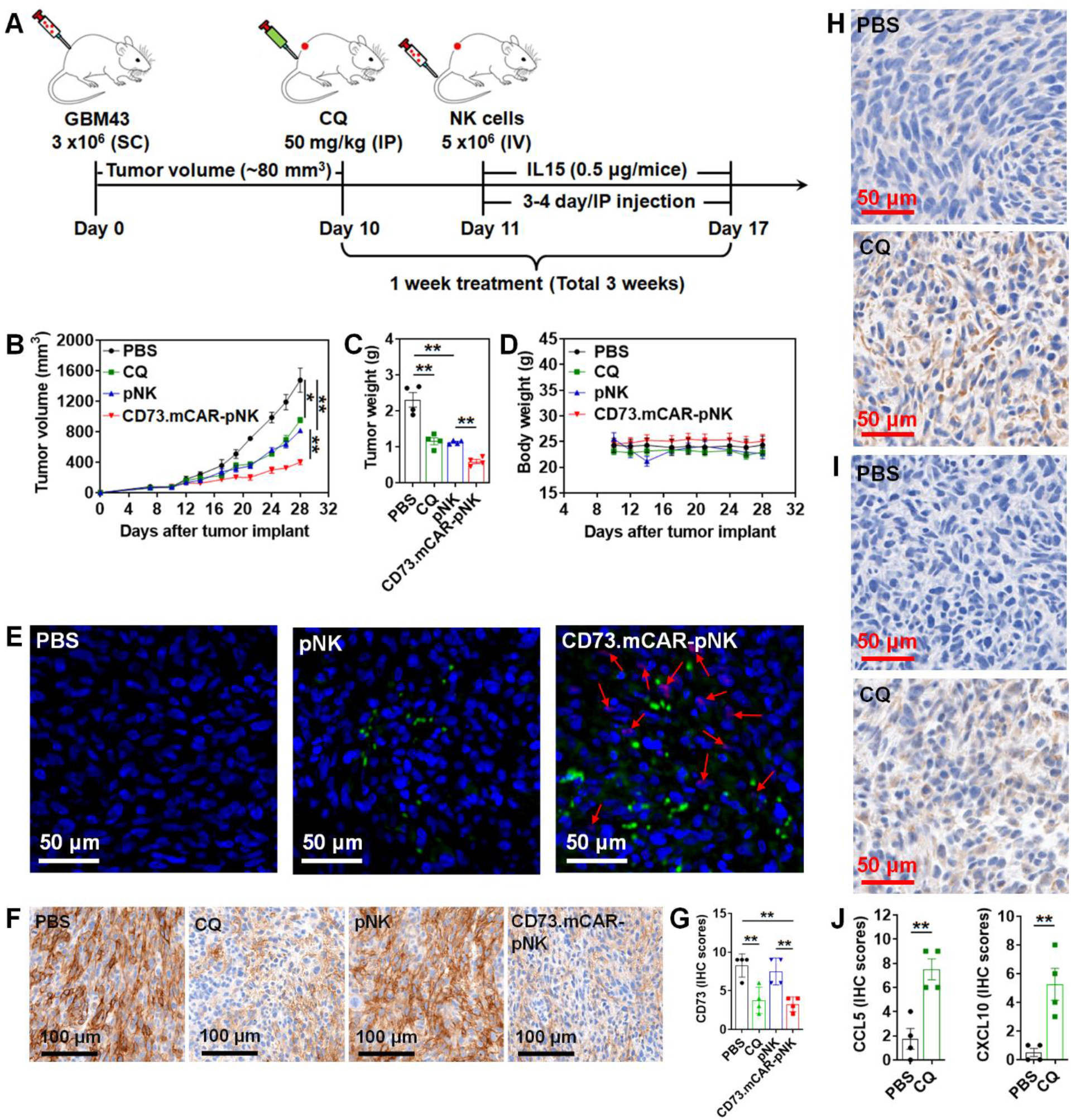
Anti-GBM activity of multifunctional engineered NK cells in a GBM43 xenograft model. (**A**) Schematic diagram illustrating the *in vivo* treatment program. (**B**) Tumor growth of individual treatment groups, including PBS, CQ, pNK cells and CD73.mCAR-pNK cells. Tumor size was determined by caliper measurements. (**C**) Average tumor weight of the mice in each treatment group after necropsy on day 28 post tumor implantation. (**D**) Changes in the body weight of the mice in each group during the treatment period. (**E**) Immunofluorescence (IF) staining of NK cells (green) and cleaved anti-CD73 scFv (arrows; red) performed on indicated tumor sections in each treatment group using anti-NKp46 antibody and Protein L. Bar = 50 μm; 200× magnification. (**F**) Immunohistochemical (IHC) staining of CD73 performed on indicated tumor sections using anti-CD73 antibody in different treatment groups. Bar = 100 μm; 200× magnification. (**G**) IHC scores of CD73 expression performed on indicated tumor sections in different treatment groups. (**H** and **I**) Immunohistochemical (IHC) staining of CCL5 (upper two panels) and CXCL10 (lower two panels) performed on indicated tumor sections in different treatment groups using anti-CCL5 and anti-CXCL10 antibodies, respectively. Bar = 50 μm; 200× magnification. (**J**) IHC scores of CCL5 or CXCL10 performed on indicated tumor sections in different treatment groups. Data are shown as mean ± SEM. \**p*<0.05, \*\**p*< 0.01.

NK cell infiltration into GBM xenograft tumors was investigated by immunofluorescence (IF) staining. Adoptively transferred NK cells (green dots) were observed in tumor tissues from pNK cell-treated mice (**Figure 5E**). In comparison, higher NK cell numbers were detected in tumors from CD73.mCAR-pNK cell-treated mice. In addition, the locally-cleaved anti-CD73 scFv (arrows; red dots) that are able to bind CD73^+^ tumor cells *in vivo* were detected in CD73.mCAR-pNK cell-treated mice in the vicinity of tumor cells. *In vitro* data reported here and in previous *in vivo* studies^52^ demonstrated the need for intratumoral protease activity to trigger ligand cleavage. For that reason, we do not expect its activation in blood and, as a result, any detectable presence of scFv in the circulation. Therefore, we used IF to detect tumor-specific presence of anti-CD73 scFv. Presence of anti-CD73 scFv co-localized only with CD73.mCAR-pNK staining, indicating the anti-CD73 scFv was successfully cleaved. Moreover, we observed a significant decrease in expression of CD73 on tumors in the CD73.mCAR-pNK treatment group compared to other treatment groups (**Figure 5F** and **G**). Additionally, tumor tissues from CQ-treated mice revealed a significantly higher presence of the chemokines CCL5 and CXCL10 (**Figure 5H-J)**. These tumors also displayed a lower level of CD73 expression (**Figure 5F** and **G**).

### Activity of the combination of multifunctional genetically-engineered NK cells with CQ against orthotopic patient-derived GBM xenografts

To further address the synergistic effect achieved by combination of CQ with CD73.mCAR-pNK cells, we established a GBM43 xenograft orthotopic model in NSG mice. Firstly, we genetically manipulated GBM43 cells to express firefly luciferase (GBM43-Luc) to enable the monitoring of tumor growth *via in vivo* bioluminescence imaging (**Figure S11**). The treatment program is shown in **Figure 6A**. Briefly, NSG mice were orthotopically implanted with GBM43-Luc cells (8 × 10^4^) and treated on day 10 post-implantation with weekly intraperitoneal (IP) injections of CQ for three weeks (50 mg/kg per day, 3 continuous days) and/or intracranial (IC) injections of CD73.mCAR-pNK cells (2 × 10^4^). When compared with those in the control group, mice treated with CQ alone showed no significant effect on tumor growth (**Figure 6B-D**). Conversely, mice that were treated with either CD73.mCAR-pNK cells or CQ + CD73.mCAR-pNK cells showed an obviously reduced tumor growth as determined by bioluminescence imaging. The most potent anti-tumor response was seen with CQ + CD73.mCAR-pNK cells treated-mice. In particular, tumors on half of the mice treated with CQ + CD73.mCAR-pNK cells showed significant arrest during the treatment period. No significant change in body weight of the mice in all groups were found throughout the entire treatment period (**Figure 6E**). Similar to subcutaneous xenograft studies, tumor tissues of mice in CQ-treated groups displayed robustly up-regulated chemokine expression, including CCL5 and CXCL10 (**Figure 6F**), which may contribute to the increased NK cell infiltration an, lead to improved therapeutic efficacy. As shown in **Figure 6G** and **H**, a significantly higher NK cell infiltration was found in GBM tumors of CQ + CD73.mCAR-pNK cells treated mice compared to mice which received CD73.mCAR-pNK cells alone. Compared to control or CQ-treated groups, the tumors of mice in both CD73.mCAR-pNK cells- and CQ + CD73.mCAR-pNK cells-treated groups displayed decreased CD73 expression, with the latter group showing the most substantial loss of CD73 expression (**Figure 6I** and **J**). This was associated with a significantly decreased level of extracellular adenosine detected in the tumors of treated mice. Specifically, CD73-mediated adenosine production in local brain tissues was decreased most prominently in mice treated with CQ + CD73.mCAR-pNK cells (**Figure 6K** and **L**).

**Figure 6.**
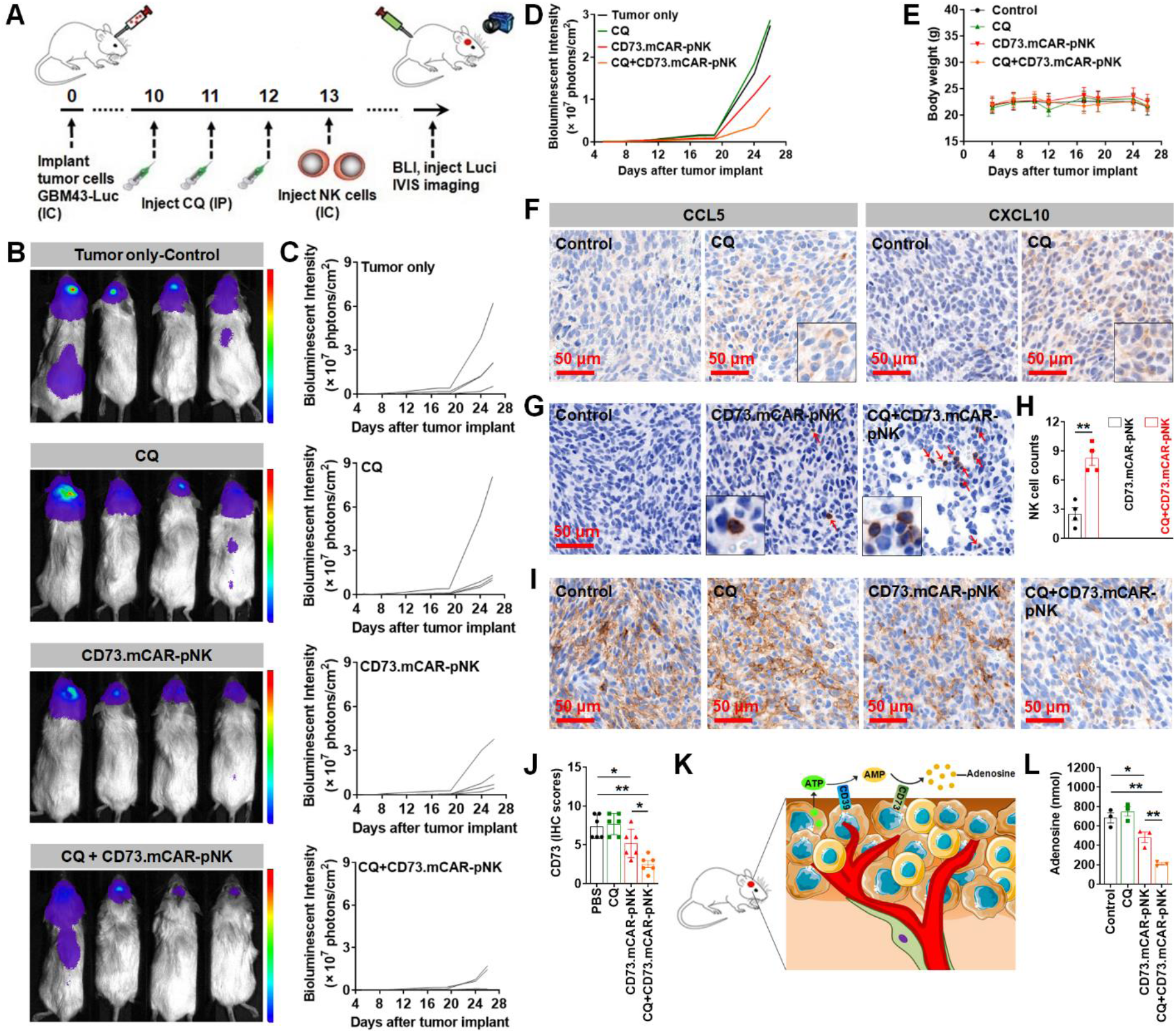
*In vivo* activity of CD73.mCAR-pNK cells in combination with the inhibition of autophagy in an orthotopic GBM43 xenograft model. (**A**) Schematic diagram showing the *in vivo* treatment program. (**B**) Bioluminescence imaging of mice in each treatment groups on day 26 after tumor implantation. (**C**) Tumor growth is displayed for individual mice of each group over time monitored using bioluminescent imaging. (**D**) Average tumor size of the mice in each group during the treatment period. (**E**) Changes in the body weight of the mice in each group during the treatment period. (**F**) Immunohistochemical (IHC) staining of CCL5 (*left two panels*) and CXCL10 (*right two panels*) performed on indicated tumor sections in different treatment groups using anti-CCL5 and anti-CXCL10 antibodies, respectively. Bar = 50 μm; 200× magnification. (**G**) Immunohistochemical (IHC) staining of NK cells performed on indicated tumor sections using anti-NKp46 antibody. Bar = 50 μm; 200× magnification. (**H**) Quantification of NK cell infiltration into intracranial tumors treated with CD73.mCAR-pNK or CQ + CD73.mCAR-pNK. Here, cell counts were recorded in 4 consecutive high-power fields (HPFs) at 200× magnification. (**I**) Immunohistochemical (IHC) staining for CD73 performed on indicated tumor sections using anti-CD73 antibody in different treatment groups. Bar = 50 μm; 200× magnification. (**J**) IHC scores of CD73 expression performed on indicated tumor sections in different treatment groups. (**K**) Schematic diagram showing extracellular adenosine production in the GBM tumor microenvironment. (**L**) Adenosine concentration in local brain tissues of mice in each treated group. Data are shown as mean ± SEM. \**p*<0.05, \*\**p*< 0.01.

Taken together, these *in vivo* xenograft studies demonstrated potent and specific activity of CD73.mCAR-pNK cells against patient-derived adult GBM. These responses were associated with the induction of significant changes in chemokine secretion and CD73 expression on GBM following co-therapy with CQ, indicating a reorganization of the GBM TME induced upon disabling autophagy.

## Discussion

GBM is a particularly challenging tumor to treat due to its intratumor heterogeneity, whereby cells with varying molecular, genetic and epigenetic characteristics contribute to its ability to evade standard treatments. This complexity results in the need for varying therapeutic strategies which can act in concert or synergy to mount meaningful therapeutic responses. The limitations of single antigen-based therapies have been recognized in clinical studies through tumor outgrowth variants. The stringent requirements for target selection^53^ place constraints on the targeting strategies that are able to result in effective and sustained responses. While combinatorial antigenic targeting with multi-targeted CAR-engineered cells has demonstrated ability to address antigen escape in GBM, poor immune cell infiltration and immunometabolic reprogramming caused by the primarily hypoxic GBM microenvironment has stymied durable responses. The unsustainable response has resulted in a lack of improvement in the OS of GBM patients treated with CAR-T therapies in Phase III trials^12^. In the current study, we describe the first example of a new approach wherein we simultaneously addressed, for the first time, three critical hurdles associated with resistance to immunotherapy for GBM: antigen escape, poor immune cell infiltration and hypoxic metabolic reprogramming. We developed an approach that strategically combines bifunctional CARs targeting GD2 along with NKG2DL and the tumor-responsive, locally-triggered release of CD73-blocking antibody fragments into a single gene-modified NK-cell product. In order to address insufficient homing of NK cells into the tumor bed, we revealed a potent reorganization of the GBM milieu induced by disabling autophagy. This resulted in substantially enhanced NK cell infiltration by engaging the CCL5 and CXCL10 chemokine axes.

We based our target selection on the combinatorial presence of pro-tumorigenic antigens in GBM patient samples. We found that gene level transcriptional expression of GD2, NKG2DL and CD73—markers with distinct roles contributing to GBM pathology―is present, individually, on over 96% (151 of 156) of the GBM TCGA patient cohort, while dual combinations can be detected on 68% of patient samples. Patient-derived GBM cells—pediatric, adult primary and adult recurrent—further confirmed this expression pattern. Modulating the strength of antigen recognition to target antigens present in low amounts can enhance activation signals in settings of poor antigenic density, but it also potentiates off-target toxicities for those antigens that are present on normal cells. Targeting a combination of antigens that couples direct antigen recognition (*via* CAR-based activation) with the sustained, spatially-controlled release of soluble antibody fragments subdues potential off-target effects and provides targeting combinations in patients for which these antigens show altered expression over time or in individual tumors. Accordingly, we armed NK cells to express a trifunctional construct which can target heterogeneous combinations of antigens responsively to their expression levels, locally and while sparing healthy cells. Specifically, these multifunctionally-armed CAR-NK cells revealed no obvious toxicity towards to normal cells in the brain, including human cerebral microvascular endothelial cells (hCMEC/D3) and human cortical neuronal cells (HCN-2). Part of this is aided by the low expression of at least one of the targeted markers on these cells. The triggerable release of CD73 antibody fragments by GBM tumor associated proteases (TAPs) results in low concentrations of CD73 in the local TME, a sustained response that depends on CAR expression but is independent of its activation, providing two related but independent mechanisms of tumor recognition.

Several studies have demonstrated significant phenotypic similarity between blood-derived NK cells from both healthy donors and GBM patients. However, the phenotype and function of GBM tumor-resident NK cells are markedly altered and characterized by significantly lower levels of the activating receptors CD16, NKG2D, NKp30, NKp46, DNAM-1, CD2 and 2B4^39^. Mechanistically, GBM patients utilize their own immunosuppressive TME to impair infiltrated NK cell function in favor of GBM escape from immune surveillance and NK-mediated cytotoxicity^54^. These immune escape mechanisms are rooted in GBM’s heterogeneity, and represent a complex network of pathways that promote GBM progression^55^. NKG2D-NKG2DL interactions play a vital role in activating the anticancer immune response^56^. However, NKG2D expression on the surface of NK cells is significantly decreased in response to the high extracellular concentrations of adenosine produced by the ectoenzyme CD73 in the GBM TME. In that context, our group has previously demonstrated that CD73 is a significant prognostic biomarker for GBM and can be used as correlative factor of GBM patient survival^32^. In this study, we induced upregulation of NKG2D by engineering NK cells to express an NKG2D-based CAR, which we previously showed can enhance anti-tumor responses in combination with CD73 blockade^57^. Immunosuppression of NK cells *via* the adenosinergic axis goes beyond activating receptor inhibition, however, and encompasses NK cell metabolism and various effector functions^36,58^. Because of the wide expression of CD73^59^, localizing targeted blockade of this enzyme may be preferential to systemic administration of anti-CD73 therapies. To achieve controlled and localized responsiveness, we engineered NK cells to possess a tumor-responsive, locally-released anti-CD73 scFv that upon triggering for release in the GBM TME is able to block local CD73 activity. Our data indicate the ability of the engineered NK cells to not only reduce the local concentration of extracellular adenosine, but also reduce the immunosuppression of NK cell activation in a tumor-specific manner. The sustained, low concentration release of anti-CD73 is likely to sustain the metabolic function of NK cells that would otherwise be lost in a setting of unencumbered CD73 enzymatic activity. In that respect, CAR-mediated signaling in combination with local CD73 blockade is likely to result in anti-tumor responses that are not only measurable by cytotoxicity against cancer cells, but also by their impaired metabolic activity, in turn sustaining the longer-term retention of NK cells in the tumor.

Temporal modulation of antigens also occurs in response to treatment. The expression of NKG2DLs, which include two MHC class I chain-related proteins (MICA and MICB) and six UL16-binding proteins (ULBP1-6), is upregulated in GBM following standard treatment with chemotherapy (TMZ) or irradiation (IR)^41^. We found NKG2DL expression to also be regulated by CQ, as treatment induced a significant increase in NKG2DL (both at MFI and % level) on patient-derived GBM cells. NKG2DL expression was previously found to inversely correlate with GBM cell maturity, with stem-like GBM cells expressing a more substantial, though heterogeneous, pattern of expression of these ligands^41,60,61^. These changes induced by CQ might sensitize GBM to effector function *via* the NKG2DL-R axis, but also promote the indirect loss of NKG2D on effector cells through a phenomenon that can be lessened by the induction of NKG2D expression *via* CARs. The reorganization of the GBM TME induced by CQ included a decrease in GD2 expression, potentially lessening the apoptotic burden imposed on tumor-infiltrating cells induced by gangliosides on GBM^62^. To recapitulate some of the stem cell-like properties associated with resistance to therapy owing to the presence of glioma stem-like cells in patient GBM tumors, we utilized patient-derived xenografts that were generated with cells sourced from patients with primary GBM.

Antigenic targeting may not be sufficient to induce sustained anti-GBM responses. Clinical data have indicated that intratumoral NK cell presence associates positively with patient outcome and survival^63,64^. However, NK cells are present in low amounts in GBM. Among regulating a variety of pathophysiological functions in GBM, chemokine receptor/ligand interactions can drive NK cell trafficking alongside their signaling gradient to result in improved homing, responses that were demonstrated for the CXCR4/CXCL12-directed NK cell homing to GBM^65^. However, scant evidence of chemokine-dependent trafficking has been shown in intracranial GBM. Our data revealed a significant relevance of the CCL5 and CXCL10 chemokine pathways in GBM upon pharmacological targeting of autophagy. Mechanistically, we found that inhibition of autophagy in GBM cells triggered the upregulation of chemokines, particularly CCL5 and CXCL10, *via* the activation of the PI3K/NF-κB signaling pathway. This subsequently conferred significant chemotactic ability to NK cells resulting in pronounced detection of adoptively-transferred CD73.mCAR-pNK cells in the brains of treated mice. Although we detected significant upregulation of these chemokines, we cannot rule out contributions from others to the intratumoral trafficking of NK cells. Conversely, expression of CCL2 and CXCL12, both recognized for their contributing roles to the recruitment of immunosuppressive cells such as tumor associated macrophages (TAMs), was significantly decreased in response to autophagy blockade^66,67^. Our preclinical *in vivo* data further demonstrated that CQ-mediated targeting of autophagy potentiates anti-GBM activity of NK cells *via* mechanisms that may involve a sensitization to their killing by modulating antigen level expression, and the activation of pro-apoptotic pathways on GBM. These data point to novel actionable responses induced by the inhibition of autophagy and CAR-based antigenic targeting. Our data also highlight the complex and sophisticated reorganization of the GBM TME induced by CQ. Confounding factors of the dichotomous effects of CQ on GBM pathology have been recognized clinically. As an adjuvant, CQ was shown to enhance clinical responses to anti-GBM therapy in double-blinded Phase III studies^68^. CQ was shown to be able to cross the blood-brain barrier *in vivo*^69–71^, while preclinical studies have demonstrated that concentrations of 50 mg/kg are suitable for administration regimens against orthotopic glioblastoma xenografts^72–74^. The multiple effects of CQ on the phenotypic signatures and chemokine profiles of GBM *in vivo*, alongside a limited ability to control tumor burden alone, however, point to a nuanced role in combination regimens which can nonetheless uniquely enhance CAR-based adoptive transfer immunotherapy. Any potential toxicity of CQ, however, needs to be factored into the therapy administration regimens^75–77^. The use of the less toxic metabolite hydroxychloroquine is one strategy to mitigate toxicity associated with CQ^78^. Intracranial injections of NK cells with lower concentrations of CQ can also be pursued. In this study, in cases where mice were to be treated with a combination of CQ and NK cells, we infused NK cells intracranially, while CQ was administered systemically. This resulted in no observable CQ-dependent toxicity in the treated groups and enhanced the effective presence of CAR-NK cells in the tumor. In this study, we showed that repeated treatments are tolerated and result in a rapid onset of response. Intracranial infusions of engineered cellular therapies have been favored over systemic ones due to the ability to achieve higher concentrations of therapeutic product locally near the tumor, and have been shown effective in humans^9^.

This study demonstrated that development of multifunctional genetically-engineered human NK (CD73.mCAR-pNK) cells can result in effective anti-GBM activity supported by a concerted approach of overcoming tumor heterogeneity and multiple immunosuppressive features of the GBM TME. We also unveiled that targeting autophagy functions as an immuno-modulator to promote the homing of effector CAR-NK cells into GBM tumor sites while reprogramming the GBM TME toward sensitization to CAR-based targeting. All treated mice experienced either a delay or complete arrest of tumor growth. The most significant responses were obtained upon co-administration of CAR-NK cells with CQ. As a first demonstration of the use of human CAR-NK cells in immunotherapy of glioblastoma, we show that while human CAR-NK therapy is a viable option, optimal responses rely on administration regiment, dosage and frequency of adoptive transfer. An issue of great interest that remains to be addressed is to evaluate the effects of targeting autophagy on the immune landscape of GBM tumors and on the infiltration of other types of immune cells into the tumor bed, which could be explored in appropriate immunocompetent models of GBM. Longer-term efficacy studies that focus on survival will be pursued in future investigations as we advance this immunotherapy toward human trials. Nevertheless, it is our belief that the findings from this study will open a new avenue and pave the way to fully exploit NK cell-based GBM immunotherapy in clinical settings.

## Methods

### Mice

Female 6- to 8-week-old Rag1^−/−^ mice and NOD.Cg-Prkdc^scid^ IL2rg^tm1Wjl^/SzJ (NSG) mice were maintained at the Purdue Center for Cancer Research. All the animal experiments described in this study were approved by the Purdue University Animal Care and Use Committee.

### Ethics Statement

Written informed consent was obtained from all subjects involved in the study. All procedures performed in studies involving human participants were approved by Purdue University’s Institutional Review Board (IRB). All institutional safety and biosecurity procedures were adhered to.

### Isolation of peripheral blood NK cells and cell culture

Primary NK (pNK) cells used in this study were obtained using Purdue University's Institutional Review Board (IRB)-approved consent forms (IRB-approved protocol #1804020540). Blood samples were obtained from healthy adult donors. And pNK cells were isolated from whole blood by negative selection using the EasySep™ Direct Human NK cell Isolation Kit (StemCell Technologies). Following isolation, the cells were cultured and expanded using NK MACS^®^ medium (130-114-429, Miltenyi) system according to manufacturer’s instructions. NK-92 cells (directly purchased from ATCC) were maintained in RPMI1640 supplemented with 10% FBS, 100 U/mL penicillin, 100 μg/mL streptomycin, 2 mM L-glutamine, 400 U/mL IL-2 and 0.1 mM 2-mercaptoethanol. HCN-2 cells (directly purchased from ATCC) were maintained in DMEM supplemented with 10% FBS, 100 U/mL penicillin and 100 μg/mL streptomycin. hCMEC/D3 cells (kindly provided by Dr. Gregory T. Knipp, Purdue University) were grown in EBM-2 supplemented with 5% FBS, 100 U/mL penicillin, 100 μg/mL streptomycin, 1.4 μM hydrocortisone, 5 μg/mL ascorbic acid, 1% chemically defined lipid concentrate, 10 mM HEPES and 1 ng/mL bFGF. SJ-GBM2, GBM43 and GBM10 cells (kindly provided by Dr. Karen E. Pollok, Indiana University School of Medicine) were grown in DMEM supplemented with 10% FBS and 1% HEPES. The SJ-GBM2 cells were originally obtained by Dr. Pollok from the Children’s Oncology Group. The GBM43 and GBM10 xenograft tissue was kindly provided by Dr. Jann Sarkaria (Mayo Clinic, Rochester MN)^79,80^. Cell lines derived from the xenograft tissue were established in Dr. Pollok’s lab according to described protocols^80^. Cell line identity was confirmed by DNA fingerprint analysis (IDEXX BioResearch) for species and baseline short-tandem repeat analysis testing as described, and were found to be 100% human^81^. All cell lines were incubated at 37 °C in a humidified 5% CO_2_ environment.

### Chemical regents

Chloroquine diphosphate salt (CQ) (98%) and adenosine 5’-monophosphate sodium salt hydrate (AMP) (99%) were purchased from ACROS Organics™. Hydrocortisone (98%) was purchased from Alfa Aesar. Ascorbic acid, Sodium chloride (NaCl) and Bovine serum albumin (BSA) were purchased from Thermo Fisher Scientific. Potassium chloride (KCl), Magnesium chloride (MgCl_2_), Sodium bicarbonate (NaHCO_3_), Adenosine (≥99%), DEAE-Dextran hydrochloride and Glucose were purchased from Sigma. Brefeldin A Solution (1000×) and Monensin Solution (1000×) were purchased from BioLegend. D-Luciferin Potassium Salt (>99%) was purchased from Syd Labs. LY294002, BAY11-782 and SP600125 were purchased from Cayman Chemical. Fetal bovine serum (FBS) and dimethyl sulfoxide (DMSO) were purchased from Corning. Recombinant human interleukin-12 (IL-2) was gifted from Akron Biotech. Recombinant human interleukin-15 (IL-15) and Fibroblast Growth Factor-basic (bFGF) were purchased from GoldBio. Recombinant Human RANTES (CCL5) was purchased from PeproTech. Recombinant human CXCL10, Recombinant human u-Plasminogen Activator/Urokinase, CF (uPA) and Recombinant human NKG2D/CD314 Fc Chimera were purchased from R&D systems. Biotin-Protein L was purchased from GenScript. BsiWI was purchased from New England Biolabs. RPMI1640, DMEM, IMDM, penicillin/streptomycin solution 100× (PS), 2-mercaptoethanol (50 mM), HEPES (1 M), chemically defined lipid concentrate, trypan blue solution and trypsin-EDTA were purchased from Gibco. Opti-MEM Reduced Serum Media was purchased from Invitrogen. EBM-2 was purchased from Lonza. Human AB serum was purchased from Valley Biomedical. Collagen I, rat tail was purchased from Enzo Life Sciences, Inc. RIPA lysis buffer system (sc-24948) was purchased from Santa Cruz Biotechnology.

### Plasmid construction and lentivirus production

(1) (2) (3) and (4) were custom-cloned and produced by vectorbuilder.com. (5) was purchased from Santa Cruz Biotechnology. (6) was purchased from GenTarget Inc.

#### (1) pT7[mRNA]-NKG2D-CAR

In this plasmid, a human NKG2D-specific CAR (NKG2D-DAP10-CD3ζ) was expressed under the control of the T7 promoter. The NKG2D sequence was derived from previous work^57^.

#### (2) pT7[mRNA]-GD2-CAR

In this plasmid, a human GD2-specific CAR (anti-GD2 scFv-CD8 Hinge-hCD28-CD3ζ) was expressed under the control of the T7 promoter. The anti-GD2 scFv sequence was derived from previous work^82^.

#### (3) pT7[mRNA]-CD73-CAR

In this plasmid, the entire construct, including a human CD73-specific cleavable anti-CD73 scFv, the human GD2-specific CAR (GD2 scFv-CD8 Hinge-hCD28-CD3ζ) and the human NKG2D-specific CAR (NKG2D-DAP10-CD3ζ) were expressed under the control of the T7 promoter. Here, the anti-CD73 scFv fragment was linked together with GD2-CAR through a (GGGGS)_3_ linker, a cleavable peptide fragment (LSGRSDNH) and a short spacer (GSSGT). The GD2-CAR was associated with NKG2D-CAR through “self-cleaving” P2A peptides (GSGATNFSLLKQAGDVEENPGP). The anti-CD73 scFv sequence was derived from previous work^83^.

#### (4) pLV[Exp]-Puro-EF1A>{CD73-CAR}

The lentiviral vector encoding the entire construct, described in (3) was expressed under the control of the EF1alpha promoter. Lentivirus production was also done by VectorBuilder. Briefly, for the third-generation lentivirus packaging, transfer plasmid (pLV[Exp]-Puro-EF1A>{CD73-CAR}) carrying the gene of interest (CD73-CAR) is co-transfected with the proprietary envelope plasmid encoding VSV-G and packaging plasmids encoding Gag/Pol and Rev into HEK293T packaging cells. After 48 hours of incubation, the supernatant is collected and centrifuged to remove cell debris and then filtered. Lentiviral particles are subsequently concentrated with PEG. The lentivirus titer was measured by the p24 ELISA. Lentivirus was stored in HBSS buffer and was shipped on dry ice.

#### (5) Beclin 1 (*BECN1*) shRNA (h) Lentiviral Particles (sc-29797-V)

*BECN1* gene knockdown *BECN1*^−^ GBM43 cells were generated by lentiviral transduction according to the manufacturer’s protocol.

#### (6) CMV-Luciferase (firefly), (Puro) Lentiviral Particles (LVP325)

The firefly luciferase-labeled GBM43 (GBM43-Luc) cells were generated by lentiviral transduction with a luciferase-bearing lentiviral vector according to the manufacturer’s protocol.

### Antibodies and stains

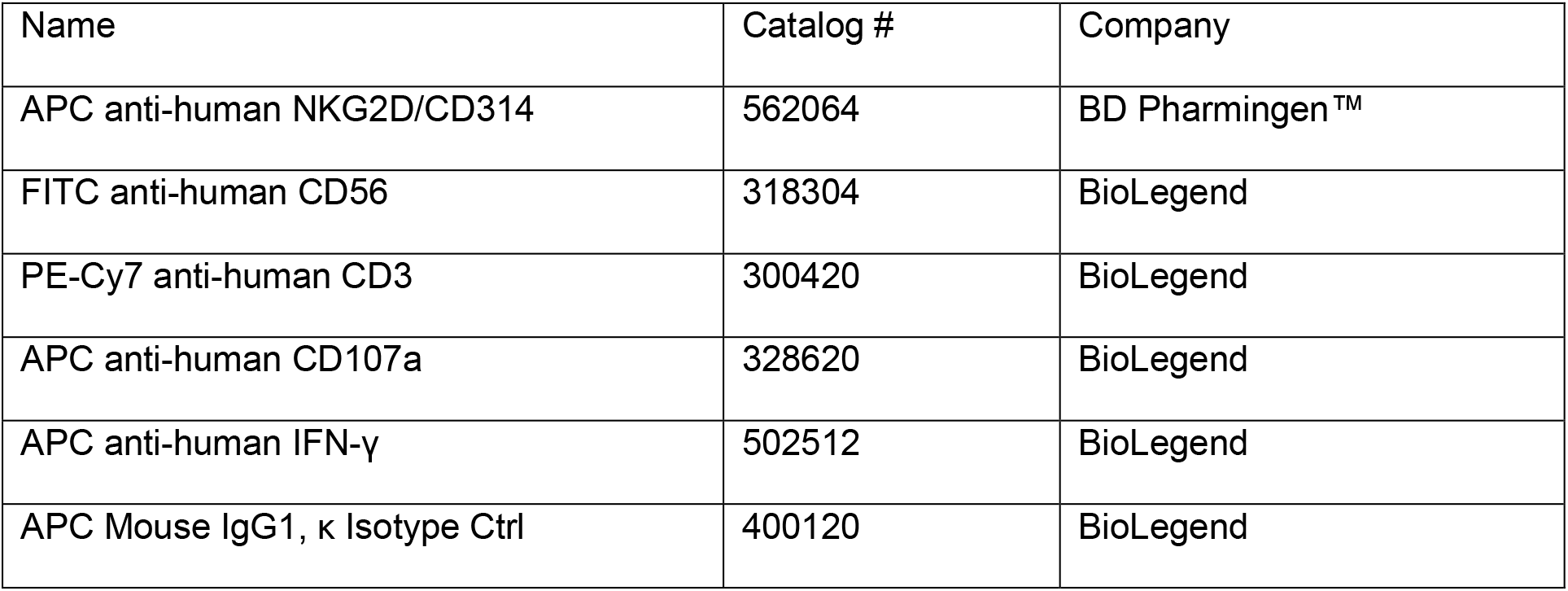
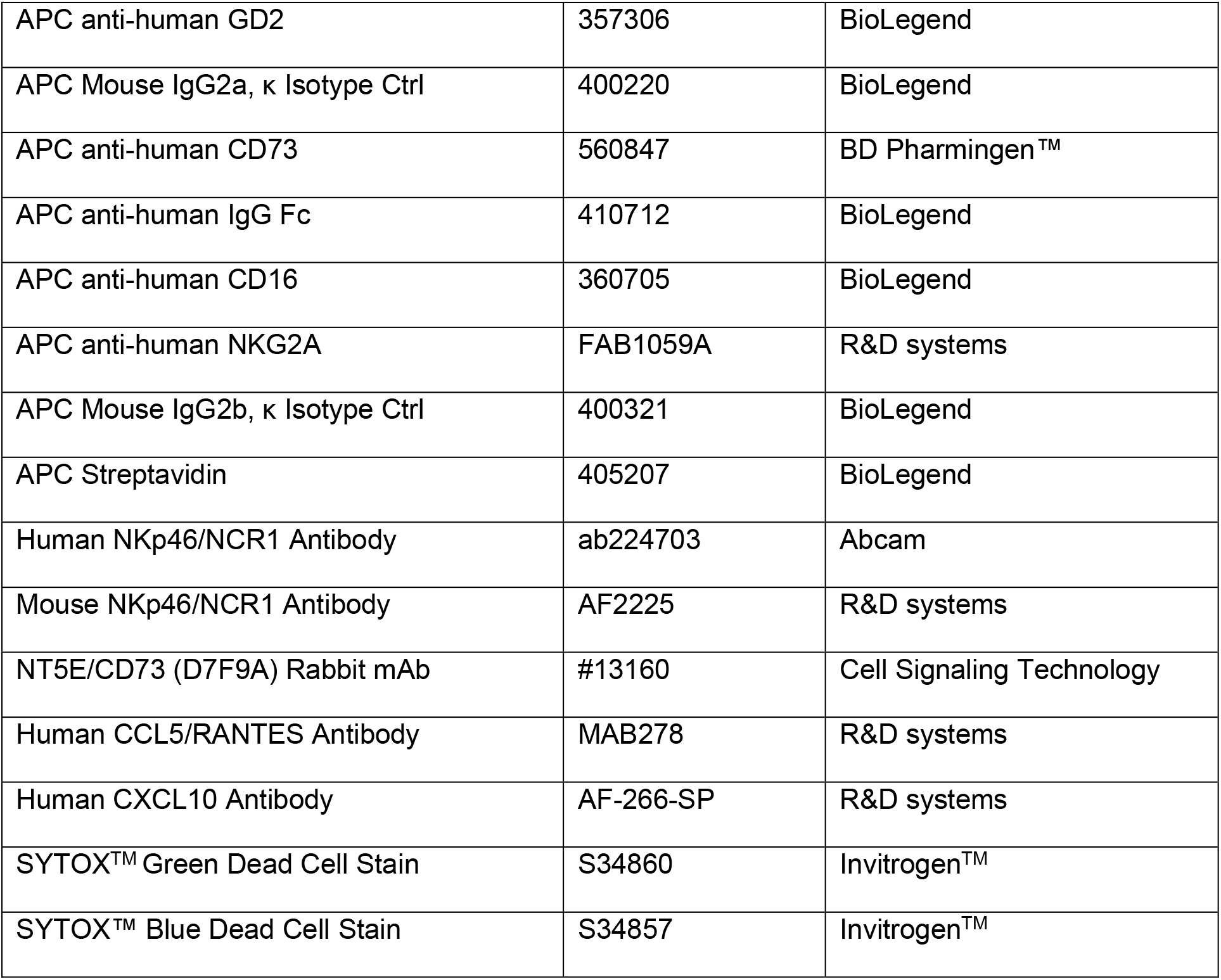

| Name | Catalog # | Company |
| --- | --- | --- |
| APC anti-human NKG2D/CD314 | 562064 | BD Pharmingen™ |
| FITC anti-human CD56 | 318304 | BioLegend |
| PE-Cy7 anti-human CD3 | 300420 | BioLegend |
| APC anti-human CD107a | 328620 | BioLegend |
| APC anti-human IFN-γ | 502512 | BioLegend |
| APC Mouse IgG1, κ Isotype Ctrl | 400120 | BioLegend |
| APC anti-human GD2 | 357306 | BioLegend |
| APC Mouse IgG2a, κ Isotype Ctrl | 400220 | BioLegend |
| APC anti-human CD73 | 560847 | BD Pharmingen™ |
| APC anti-human IgG Fc | 410712 | BioLegend |
| APC anti-human CD16 | 360705 | BioLegend |
| APC anti-human NKG2A | FAB1059A | R&D systems |
| APC Mouse IgG2b, κ Isotype Ctrl | 400321 | BioLegend |
| APC Streptavidin | 405207 | BioLegend |
| Human NKp46/NCR1 Antibody | ab224703 | Abcam |
| Mouse NKp46/NCR1 Antibody | AF2225 | R&D systems |
| NT5E/CD73 (D7F9A) Rabbit mAb | #13160 | Cell Signaling Technology |
| Human CCL5/RANTES Antibody | MAB278 | R&D systems |
| Human CXCL10 Antibody | AF-266-SP | R&D systems |
| SYTOX™ Green Dead Cell Stain | S34860 | Invitrogen™ |
| SYTOX™ Blue Dead Cell Stain | S34857 | Invitrogen™ |

### Assay Kits

1. TransIT^®^-mRNA Transfection Kit (Mirus Bio LLC);
2. EasySep™ Direct Human NK cell Isolation Kit (StemCell Technologies);
3. Human CCL5 (RANTES) Biolegend-ELISA MAX^TM^ Deluxe Sets (BioLegend);
4. Human CXCL10 (IP10) Biolegend-ELISA MAX^TM^ Deluxe Sets (BioLegend);
5. Malachite Green Phosphate Assay Kit (BioAssay Systems);
6. Adenosine Assay Kit (Cell Biolabs Inc.);
7. mirVana^™^ miRNA Isolation Kit (Life Technologies);
8. HiScribe^™^ T7 ARCA mRNA Kit (with tailing) (New England Biolabs);
9. EZ-10 Spin Column RNA Cleanup &Concentration Kit (Bio Basic Inc.);
10. qScript^™^ One-Step SYBR^®^ Green qRT-PCR Kit,Low ROX^™^ (Quanta BioSciences Inc.);
11. Fixation/Permeabilization Solution Kit (BD Biosciences);
12. 7-AAD/CFSE Cell-Mediated Cytotoxicity Assay Kit (Cayman Chemical).

### RT-PCR Primers

Human CCL5 qPCR primer pair (HP100784), Human CXCL10 qPCR pair (HP100690), Human CCL2 qPCR primer pair (HP104854), Human CXCL9 qPCR primer pair (HP100773) and Human CXCL12 qPCR primer pair (HP100192) were obtained from Sino Biological Inc. GAPDH was used as the endogenous control. Its primers—primer 1 (5’-ACATCGCTCAGACACCATG-3’) and primer 2 (5’-TGTAGTTGAGGTCAATGAAGGG-3’)—were obtained from Integrated DNA Technologies, Inc. All the primers described in the RT-PCR assay were used according to the manufacturer’s instructions.

### Bioinformatics analysis of GBM patient data

Glioblastoma (GBM) RNA-seq data (156 patients) was downloaded from TCGA^46^ through the Genomic Data Commons. Linear correlation between normalized expression (FPKM) of selected genes was determined using corrplot R-package. Next, the GBM Patients (N = 156) were classified into high/low groups based on expression of each individual gene (*B4GALNT1, MICA, MICB* and *NT5E*) using the upper and lower quartiles as cutoff for high and low expression, respectively. The high expression group was of interest and number of patients with high expression of at least one of the 4 genes was used to build a Venn diagram.

### mRNA synthesis and engineered NK-92 cell generation

mRNA individually derived from plasmid (1), (2) and (3) was synthesized through HiScribe^™^ T7 ARCA mRNA Kit (with tailing) (New England Biolabs). All the mRNA products were concentrated and purified *via* EZ-10 Spin Column RNA Cleanup & Concentration Kit (Bio Basic Inc.). After that, their concentrations were measured using a Qubit 4 Fluorometer (Fisher Scientific). To generate the engineered NK-92 cells with the expression of different CAR structures, NK-92 cells were transfected with mRNA *via* TransIT^®^-mRNA Transfection Kit (Mirus Bio LLC) according to the manufacturer’s recommended protocols. Briefly, NK-92 cells were plated in 12-well plates (5 × 10^5^ cells/well in 1 mL complete growth medium). Then, the mRNA (1 μg) was added into Opti-MEM (100 μL) in a sterile polystyrene tube and mixed well. After that, the mRNA Boost Reagent (3 μL) and Trans Reagent (3 μL) were added and mixed sequentially. The tube was incubated at room temperature for 5 min, and the complexes were added dropwise to the cells. Following transfection, the cells were incubated at 37°C and 5% CO_2_ for 48 hours and then harvested for the CAR structures expression determination. To measure the transfection efficiency *via* the levels of NKG2D expression, the cells were stained with APC-conjugated NKG2D antibody and analyzed by flow cytometry using a BD Accuri™ C6 Plus (Becton Dickinson). For detection of GD2.CAR expression, the cells were sequentially stained with Biotin-Protein L and APC-streptavidin. Then, the levels of GD2.CAR expression were analyzed by flow cytometry. For CD73.mCAR-NK-92 cells expressing the full CAR construct, expression of anti-CD73 scFv was determined out by incubating the cells with various concentrations of uPA to trigger the removal of anti-CD73 scFv from the cell surface and the rest of the CAR-construct. Following this, the cells were stained with Biotin-Protein L and APC-streptavidin. The expression of the remaining construct was then analyzed by flow cytometry. The cell viability after transfection was measured by CCK-8 (Dojindo Molecular Technologies, Inc) assay analysis.

### Generation of engineered primary NK (CD73.mCAR-pNK) cells

Cytokine-activated pNK cells (activated for 1 week of culture in NK MACS^®^ medium) were engineered to express the multifunctional CAR over two rounds of lentiviral transduction with the help of dextran. Briefly, pNK cells were plated in 24-well plates (5×10^5^ cells/well in 0.5 mL RPMI1640 medium supplemented with 10% FBS and 500 U/ml IL-2). Then, the lentivirus suspension was added at 10 multiplicity of infection (MOI). After that, the dextran aqueous solution was added to a final concentration of 8 μg/mL. Finally, the plate was centrifuged at 1000g for 60 min and incubated overnight in a 37 °C incubator infused with 5% CO_2_. After the transduction, the CD73.mCAR-pNK cells were harvested and expanded under the NK MACS medium until to further use. The expression level of all the constructs was measured and analyzed by flow cytometry according to above-described protocol. The cell viability after transduction was measured by Trypan blue staining.

### Functional *in vitro* analysis of engineered NK cells

#### Killing assay

To test for killing potency, we used a killing assay as described previously. Either NK-92, NKG2D.CAR-NK92, GD2.CAR-NK92, CD73.mCAR-NK92, pNK or CD73.mCAR-pNK cells were co-cultured with different target cells (SJ-GBM2, GBM43, GBM10, hCMEC/D3 and HCN-2) at various E:T ratios for 4 hours. To evaluate the effects of anti-CD73 scFv part on the killing activity, either CD73.mCAR-NK92 or CD73.mCAR-pNK cells with expression of the entire construct were incubated with uPA to remove all the anti-CD73 scFv from the surface. After that, the killing ability of these cells against GBM43 target cells was measured.

#### Degranulation

To test for degranulation, we followed a protocol as previously described. Briefly, either CD73.mCAR-NK92 or CD73.mCAR-pNK cells were co-cultured with GBM43 cells (E:T = 5:1) for 4 hours in the presence of APC-CD107a antibody and monensin. Detection of CD107a (% and MFI) was carried out by flow cytometry.

#### Cytokine production

To test for IFN-γ production, we followed a protocol as previously described. Briefly, either CD73.mCAR-NK92 or CD73.mCAR-pNK cells were co-cultured with GBM43 cells (E:T = 5:1) for 4 hours in the presence of brefeldin A. Then, the cells were collected, washed, and fixed/permeabilized. After that, the cells were stained with APC-IFN-γ antibody and detected by flow cytometry.

#### CD73 enzymatic activity

To test for CD73 activity and its ability to generate adenosine, GBM43 cells were seeded at 2 × 10^4^ cells per well in a 96-well plate in complete DMEM. After overnight incubation, GBM43 cells were incubated with anti-CD73 scFv following uPA (100 nM)-mediated cleavage from either CD73.mCAR-NK92 or CD73.mCAR-pNK cells during incubation for 6 hours. Then, the cell culture medium was removed and cells were rinsed three times with phosphate-free buffer (117 mM NaCl, 5.3 mM KCl, 1.8 mM MgCl_2_, 26 mM NaHCO_3_, 10 mM glucose, pH 7.4, diluted in ddH_2_O). AMP (250 μM final), diluted in phosphate-free buffer, was added and incubated for 10 min at 37 °C. Finally, the phosphate (Pi) concentrations resulting from AMP hydrolysis were measured using a malachite green phosphate assay kit following the manufacturer’s instructions.

#### Effect of CAR expression on NK cell phenotype changes in response to GBM

To test for changes in CD16 and NKG2A expression on NK cells in response to GBM cells, CFSE-labeled GBM43 cells were first seeded at 4 × 10^4^ cells per well in a 24-well plate in complete DMEM. After overnight incubation, either pNK or CD73.mCAR-pNK cells were added (E:T = 5:1) for 4 hours. Then, the cells were collected, washed and stained with either APC-CD16 or APC-NKG2A antibody and the expression was measured by flow cytometry.

### Inhibition of autophagy

#### Generation of BECN1 knockdown GBM cells (*BECN1*^−^ GBM43)

GBM43 cells were grown as previously described. Lentiviral *BECN1* shRNA particles were used, according to the manufacturer’s instructions, to generate *BECN1*^−^ GBM43 cells. *In vitro* growth behavior of *BECN1*^−^ GBM43 was verified *via* CCK-8 assay analysis.

#### Pharmacological and genetic inhibition studies

For pharmacological inhibition of autophagy, GBM43 cells were treated with different concentrations of CQ. To determine *in vitro* cell viability in response to CQ, GBM43 cells were seeded in 96-well plates at a density of 1 × 10^4^ cells/well. After overnight culture, the cells were treated with different concentrations of CQ and incubated for another 24 hours. The cell viability was then determined with CCK-8 assay analysis.

#### Western Blotting

Cell lysates were prepared using RIPA lysis buffer system according to the manufacturer’s instructions. After that, the samples were run and analyzed *via* auto-western service provided from RayBiotech, Inc. The target autophagic markers include BECN1, LC3B, p62 and β-actin.

### RNA extraction and RT-PCR

Total RNAs was extracted using the mirVana^™^ miRNA Isolation Kit and the concentrations determined with a Qubit 4 Fluorometer. The RNA (80 ng) from each sample was reverse-transcribed using the qScript^™^ One-Step SYBR^®^ Green qRT-PCR kit in a ViiA-7 RT-PCR system (Thermo Fisher Scientific). The *GAPDH* gene was used as the endogenous control. The comparative Ct values of genes of interest were normalized to the Ct value of *GAPDH*. The 2^−Δct^ method was used to determine the relative expression of the genes, while the 2^−ΔΔct^ method was used to calculate fold changes of gene expression over control.

### ELISA measurements

To measure secreted CCL5 and CXCL10, GBM43 cells were treated with CQ at a final concentration of 50 μM in presence or absence of various small molecule inhibitors, including LY294002, BAY11-7082 and SP600125 for 24 hours. Supernatants were collected and the levels of CCL5 and CXCL10 were quantified using Human CCL5 and CXCL10 Biolegend-ELISA MAX^TM^ Deluxe Sets according to the manufacturer’s directions.

### Transwell BBB migration assay

An *in vitro* direct contact co-culture blood-brain barrier (BBB) model was established as previously described^51^. Briefly, human astrocytes were seeded at a density of 4 × 10^4^ cells/cm^2^ into 24-well transwell^®^ inserts (pore size 5 μm) pre-coated with 2 μg/cm^2^ poly-L-lysine and allowed to proliferate/differentiate for 48 hours in media. Then, media were removed and hCMEC/D3 cells suspended into EBM-2 media were seeded at a density of 1 × 10^5^ cells/cm^2^. The co-culture was grown in EBM-2 with media changes every other day for an additional 7 days before studies were conducted. 600 μL of RPMI1640 medium supplemented with 1% FBS containing recombinant CCL5 or CXCL10 at 100 ng/mL was placed in the lower chamber of the transwell^®^ plate. Activated pNK cells (5 × 10^5^) in 100 μL RPMI1640 medium supplemented with 1% FBS were placed into the upper chamber (5-μm pore size). After incubation for 6 hours at 37°C and 5% CO_2_, the number of pNK cells that migrated into the lower chamber was determined by flow cytometry. Data are presented as percentage of migration based on total cell input.

### *In vivo* animal studies

#### BECN1 knockdown GBM xenograft studies

GBM43 cells (3 × 10^6^) were inoculated subcutaneously (SC) in both flanks (GBM43 control cells in the right flanks and *BECN1*^−^ GBM43 cells in the left flanks) of Rag1^−/−^ mice. The tumor growth was monitored and the length (L), width (W) and height (H) of the tumor were measured using a digital caliper. The tumor volume (mm^3^) was calculated using the formula: V = 0.52 × L× W × H. Tumor growth was monitored until mice met predefined endpoint criteria. At that point, the tumor tissues were harvested and processed for histologic (IHC) analyses.

#### Efficacy studies in subcutaneous patient-derived GBM xenografts

Subcutaneous GBM xenografts were established by inoculating NSG mice with 3 × 10^6^ GBM43 cells in the right flank. Ten days later, when tumors reached a volume of about 80 mm^3^, mice were randomly assigned to 5 groups (n = 4/group) and treated according to one of the following protocols: (1) PBS only; (2) intravenous (IV) injection of 5 × 10^6^ pNK cells alone; (3) intravenous (IV) injection of 5 × 10^6^ CD73.mCAR-pNK cells alone; (4) intraperitoneal (IP) injection of CQ (50 mg/kg) alone. NK cells were administered once a week for three weeks. The mice receiving adoptive NK cells therapy (groups 2 and 3) also received IL-15 (0.5 μg/mice) by IP injection every 3-4 days. Tumor growth was followed by caliper measurements and tumor volumes were calculated using the formula: V = 0.52 × L× W × H as described above. Body weights of the mice were also recorded during the treatment. At the end of the therapy, the mice were sacrificed and tumors were harvested for histologic (IHC/IF) analyses.

#### Immunotherapy in patient-derived intracranial GBM xenografts

Orthotopic GBM xenografts were generated using NSG mice as previously described^81^. Briefly, on day 0, the GBM43-Luc cells (1 × 10^5^) were stereotactically implanted into the right forebrain through a burr hole created using a digitalized stereotactic delivery system. 9 days after implantation, mice were randomly assigned to 4 groups (n = 4/group) and treated according to one of the following protocols: (1) untreated (as the control); (2) intraperitoneal (IP) injection of CQ (50 mg/kg) alone; (3) intracranial (IC) injection of 2 × 10^6^ CD73.mCAR-pNK cells alone (suspended in 5 μL 1% BSA (IgG-free) in PBS); (4) combination of CQ (IP) and CD73.mCAR-pNK cells (IC). CQ was continually injected 3 times a week (once/day) before administration of CD73.mCAR-pNK cells. CD73.mCAR-pNK cells were administered once a week for 2 weeks *via* direct intracranial injection. CD73.mCAR-pNK cell injections were performed using a syringe through the same burr hole. Tumor volumes were monitored and recorded using the Spectral Ami Optical imaging system. Body weights of the mice were also recorded during the treatment period. At the end of the treatment, the mice were sacrificed and whole brain tissues from each mouse were harvested for adenosine measurement and histological (IHC/IF) analyses.

### Immunohistochemistry (IHC) and Immunofluorescence (IF) staining

Immunohistochemistry (IHC) and immunofluorescence (IF) staining were carried out at the Histology Research Laboratory at the Purdue University College of Veterinary Medicine. Briefly, the tumors were fixed in 10% neutral-buffered formalin, embedded in paraffin, and cut into 3–5 μm sections.

For *BECN1*^−^ GBM43 xenografts, the mouse NK cells in tumors were detected through the staining using mouse NKp46/NCR1 antibody. For the quantification of NK cells in the tumors, the stained cells were counted in 4 randomly selected intratumoral fields of each slide at 200× magnification. And the CCL5 and CXCL10 expression in the tumors were detected through the staining using human CCL5/RANTES antibody and CXCL10 antibody respectively. The staining was evaluated based on the intensity (weak = 1, moderate = 2, and high = 3) of chemokine immunostaining and the density (0% = 0, 1-40% = 1, 41-75% = 2, >76% = 3) of positive tumor cells. The final score of each sample was multiplied by the intensity and density.

For subcutaneous GBM43 xenografts, NK cell infiltration was investigated by immunofluorescence (IF) staining, which was performed with the following stains: Protein L (Alexa 647, far red), NKp46 (Alexa 488, green) and DAPI nuclear counterstain. CCL5 and CXCL10 expression were evaluated by IHC staining as described above. CD73 expression in the tumors was detected through the staining using human CD73 antibody.

For intracranial GBM xenografts, the NK cells in tumors were detected through the staining using human NKp46/NCR1 antibody. For the quantification of NK cells in the tumors, the stained cells were counted in 4 randomly selected intratumoral fields of each slide at 200× magnification. The CCL5 and CXCL10 expression in the tumors were detected through the staining using human CCL5/RANTES antibody and CXCL10 antibody respectively. CD73 expression in the tumors was detected through the staining using human CD73 antibody. The staining was evaluated as above-described method.

### Determination of intratumor adenosine concentration

Brain tissues were harvested post-treatment, rinsed with cold PBS and homogenized in PBS. After that, the suspension was centrifuged at 10,000g for 10 minutes at 4°C and the supernatant was collected. Adenosine concentrations were determined using an Adenosine Assay Kit according to the manufacturer’s directions.

### Statistical analysis

Data were presented as mean ± SEM. Statistical analysis was performed using Excel 2007 software (Microsoft office 2007). Comparison between two normally distributed test groups was performed using the two-tailed Student’s *t*-test. For analysis of three or more groups, comparison was performed using a one-way ANOVA analysis. *p*< 0.05 was considered to be statistically significant.

## Acknowledgements

This work was supported by NIH Shared Instrumentation Grant (#S10DO20029; Flow cytometry), the V Foundation for Cancer Research (Grant #D2019-039) and the Walther Cancer Foundation (Embedding Tier I/II Grant #0186.01).

## Author contributions

J.W. and S.M. conceived, designed and executed the experiments and wrote the manuscript; B.E. and S.T-A. maintained mice and carried out *in vivo* xenograft studies; N.A.L. and S.G. performed bioinformatics analysis; V.B-C. performed histological analysis; M.B. and G.T.K. developed and provided support with *in vitro* BBB model studies; K.P., M.V., A.S. and Y.Y. helped establish *in vivo* intracranial mouse models and provided support for intracranial work; K.P. helped establish patient-derived GBM lines; K.N. and R.B. provided support with experimentation and analysis of analytical NK cell studies and GBM targeting studies; all authors reviewed and edited the manuscript.

