## Supplementary Information for "Tumor-responsive, multifunctional CAR-NK cells cooperate with impaired autophagy to infiltrate and target glioblastoma"

**Table S1**

**Figure S1-S12**

**Videos 1-2**

| Pathway | pval | padj | ES | NES | Direction |
|---|---|---|---|---|---|
| ***MICA/MICB*** | 0.1374046 | 0.1374046 | 0.7373083 | 1.351678 | Positive |
| ***NT5E*** | 0.26546 | 0.26546 | -0.6885178 | -1.184308 | Negative |
| ***B4GALNT1*** | 0.2709677 | 0.2709677 | -0.6425565 | -1.228292 | Negative |
| ***CXCL10*** | 0.0072115 | 0.0072115 | 0.9004993 | 1.752424 | Positive |
| ***CCL5*** | 0.0275424 | 0.0275424 | 0.8680867 | 1.609939 | Positive |

**Table S1.** Correlation between normalized expression (FPKM) of the entire NK five-gene set and individual genes *MICA/MICB, NT5E, B4GALNT1, CXCL10* and *CCL5* based on TCGA GBM data. NES: enrichment score; ES: normalized enrichment score.

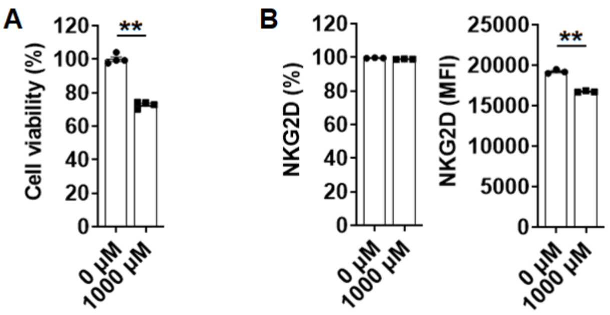


**Figure S1. Effects of adenosine on pNK cell viability and NKG2D expression.** (**A**) Cell viability (%) of pNK cells after treatment with 1000 μM of adenosine for 24 h. (**B**) NKG2D expression on pNK cells after treatment with 1000 μM of adenosine for 24 h. Note: pNK cells were seeded into 24-well plate at a density of $5 \times 10^5$ cells/well in 500 μL medium. Adenosine was then added to a final concentration of 1000 μM. After 24 h, the cells were collected and their viability (%) was determined via a CCK-8 assay. The NKG2D expression (% and MFI) was determined by flow cytometry. Data are shown as mean ± SEM.

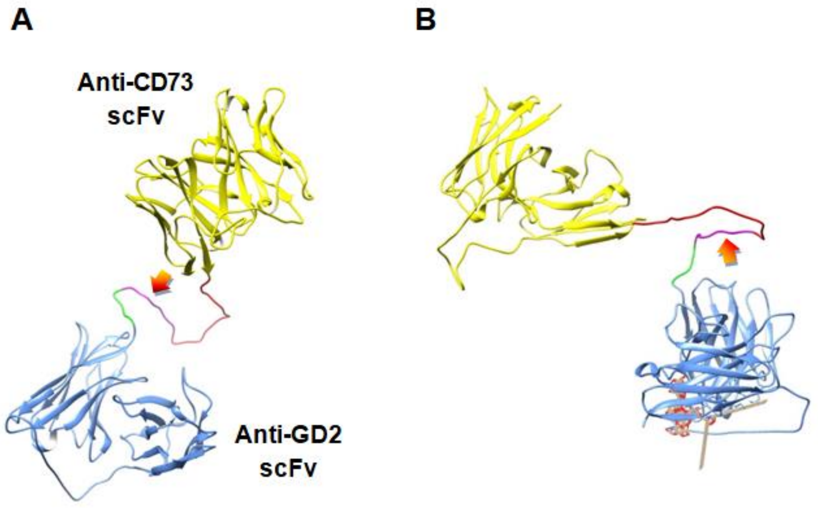


**Figure S2.** ***In silico*** **modeling of the tandem GD2.CD28.CD3ζ-CAR-CD73 scFv structure.** (**A**) Molecular model of tandem anti-GD2 (blue) and anti-CD73 scFv (yellow) extracellular domains. The model was obtained based on vector sequences using RaptorX. The cleavable peptide is shown in pink (arrows), GS spacers are shown in red and the linkers in green. (**B**) Docking of extracellular scFv domains with GD2. Docking was performed using PatchDock and refined in FireDock using a model of the entire extracellular region generated prior. **I**mages were generated in Chimera.

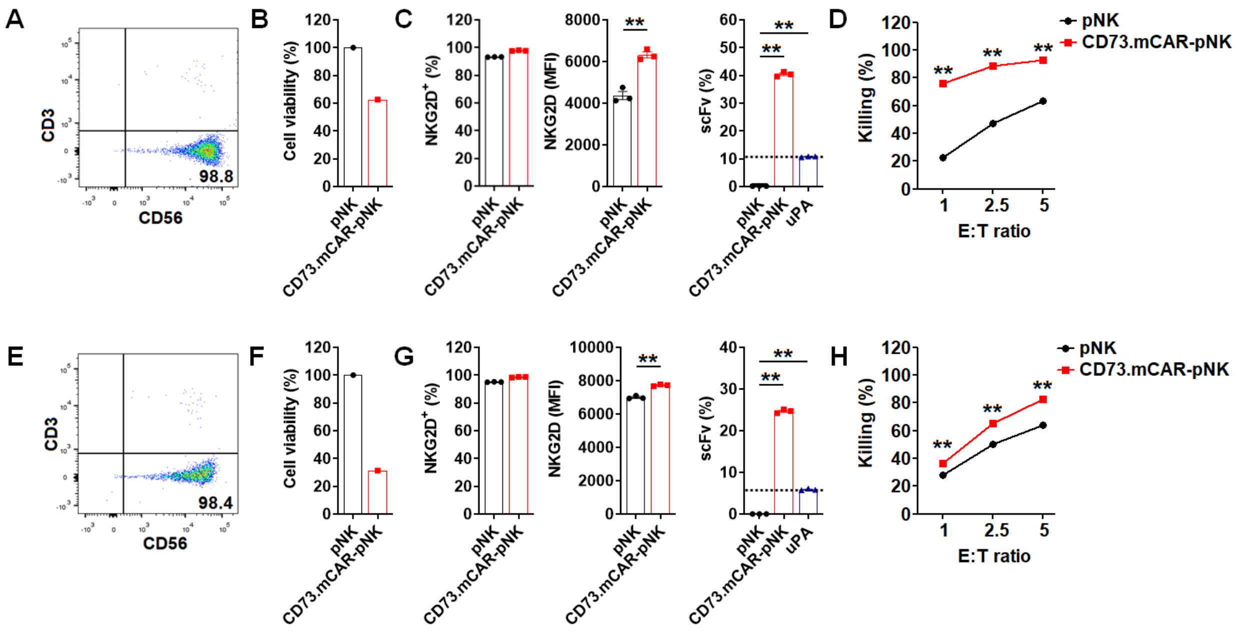


**Figure S3. Killing of patient-derived GBM43 cells by CD73.mCAR-pNK cells engineered from multiple donors.** (**A**-**D**) and (**E**-**H**) are the data for the pNK cells isolated from another two separate donors. The characterization and expression measurement were investigated as described in *Methods*. NK cells were co-incubated with GBM43 cells at different E/T ratios for 4 h and killing was quantified as described in *Methods*. Data are shown as mean ± SEM. $*p < 0.05$, $**p < 0.01$.

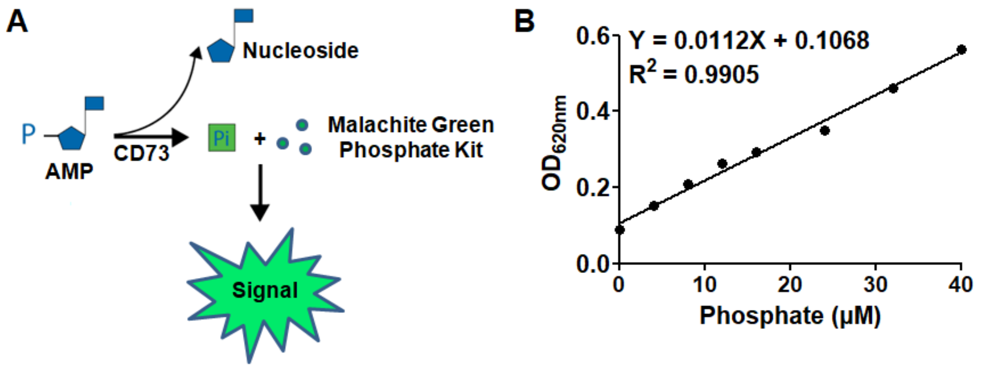


**Figure S4. Determination of CD73 enzyme activity on GBM43 cells.** (**A**)Schematic diagram of the reaction employed to detect phosphate formation by CD73. (**B**) Phosphate standard curve in 96-well plate. After a 30-minute incubation, the OD value at 620 nm was read.

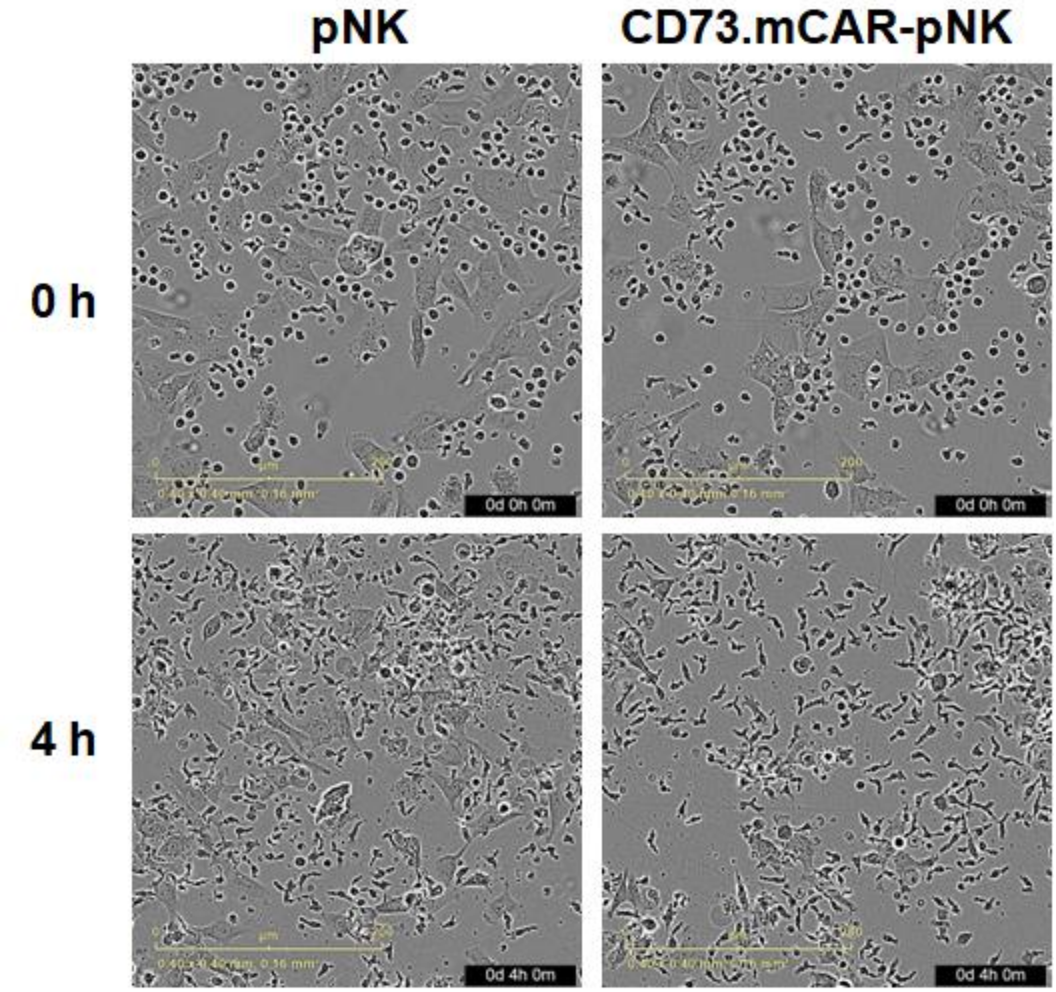


**Figure S5. Live imaging micrographs of co-culture of GBM43 cells with pNK or CD73.mCAR-pNK cells.** GBM43 cells were seeded into a 24-well plate at a density of 4 × $10^4$ cells/well. After overnight culture, either pNK or CD73.mCAR-pNK cells were added at an E/T ratio at 5. The co-cultures were then imaged using an IncuCyte S3 with scans performed every 10 min for 4 h. The small irregularly-shaped circular dots in the images represent NK cells, and the large fusiform cells represent GBM43 cells. Scale bar = 200 µm.

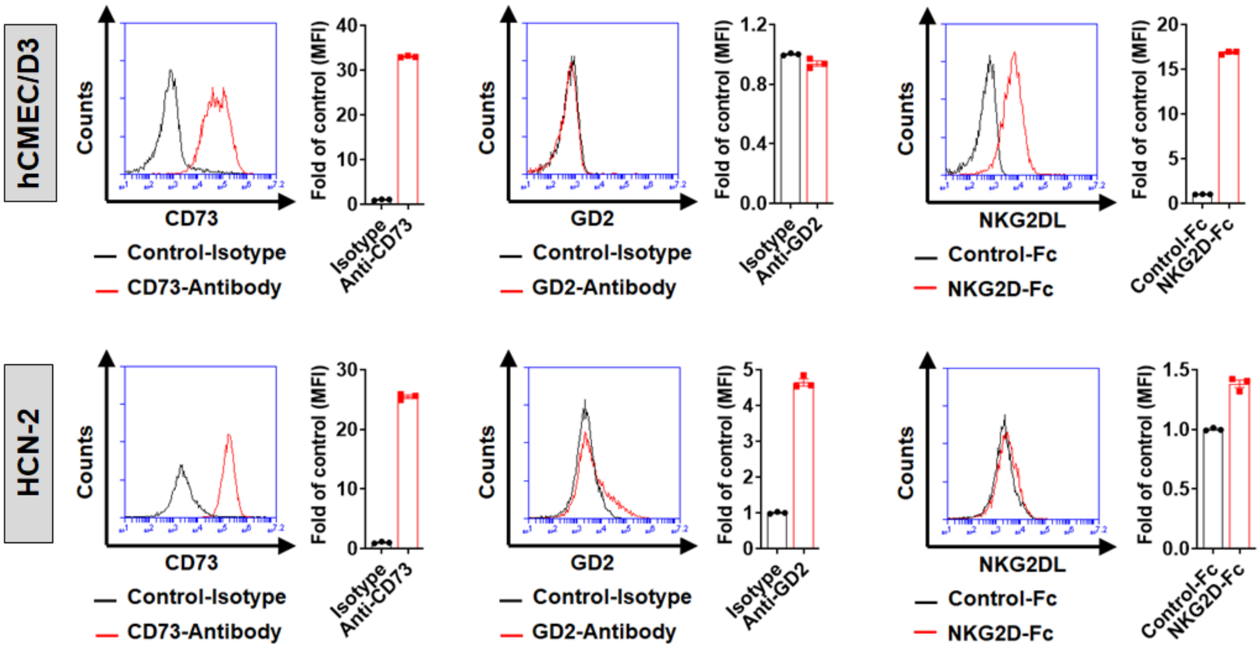


**Figure S6. Surface expression of CD73, GD2 and NKG2DL on hCMEC/D3 and HCN-2 cells.** Expression levels of the three markers on normal brain cell lines were determined by flow cytometry and are reported as fold-change (FC) over control. Data are shown as mean ± SEM.

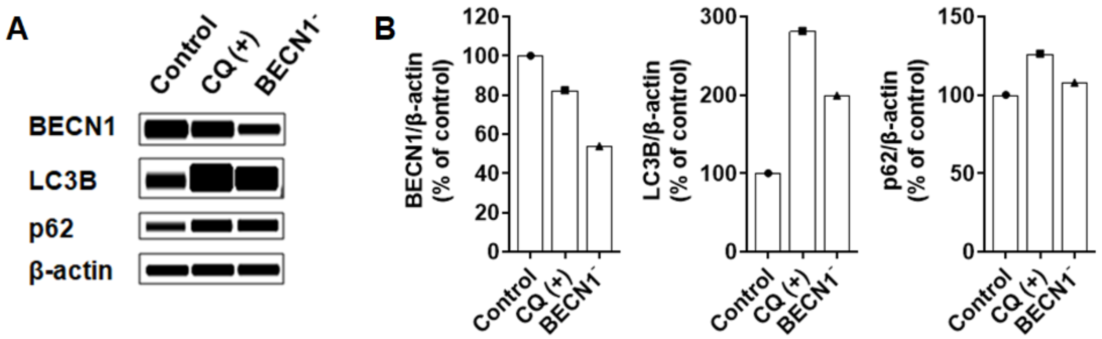


**Figure S7. Inhibition of autophagy in GBM cells either through *BECN1* gene knockdown or pharmacological treatment with CQ.** (**A**) GBM43 cells were transfected with *BECN1* shRNA (h) lentiviral particles or treated with 50 μM of CQ for 24 h. Following this, the cells were lysed and expression levels of BECN1, LC3B and p62 were analyzed via flow cytometry. β-actin was used as the loading control. (**B**) The relative expression levels of BECN1 (BECN1/β-actin), LC3B (LC3B/β-actin) and p62 (p62/β-actin) were quantified. Note: Beclin 1 (BECN1), LC3-II (LC3B) and p62 are the three primary markers of autophagy.

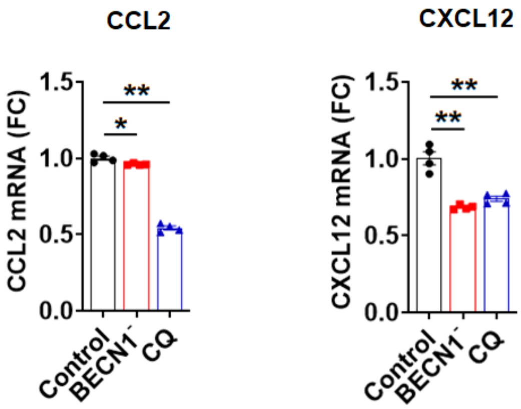


**Figure S8. Chemokine expression in response to inhibition of autophagy on GBM.** Expression of CCL2 and CXCL12 mRNA in *BECN1*$^{-}$ GBM43 or CQ-treated GBM43 cells was determined by RT-PCR. *GAPDH* was used as the reference gene. Results are reported as a fold-change (FC) over control. Data are shown as mean ± SEM. *$p < 0.05$, **$p < 0.01$.

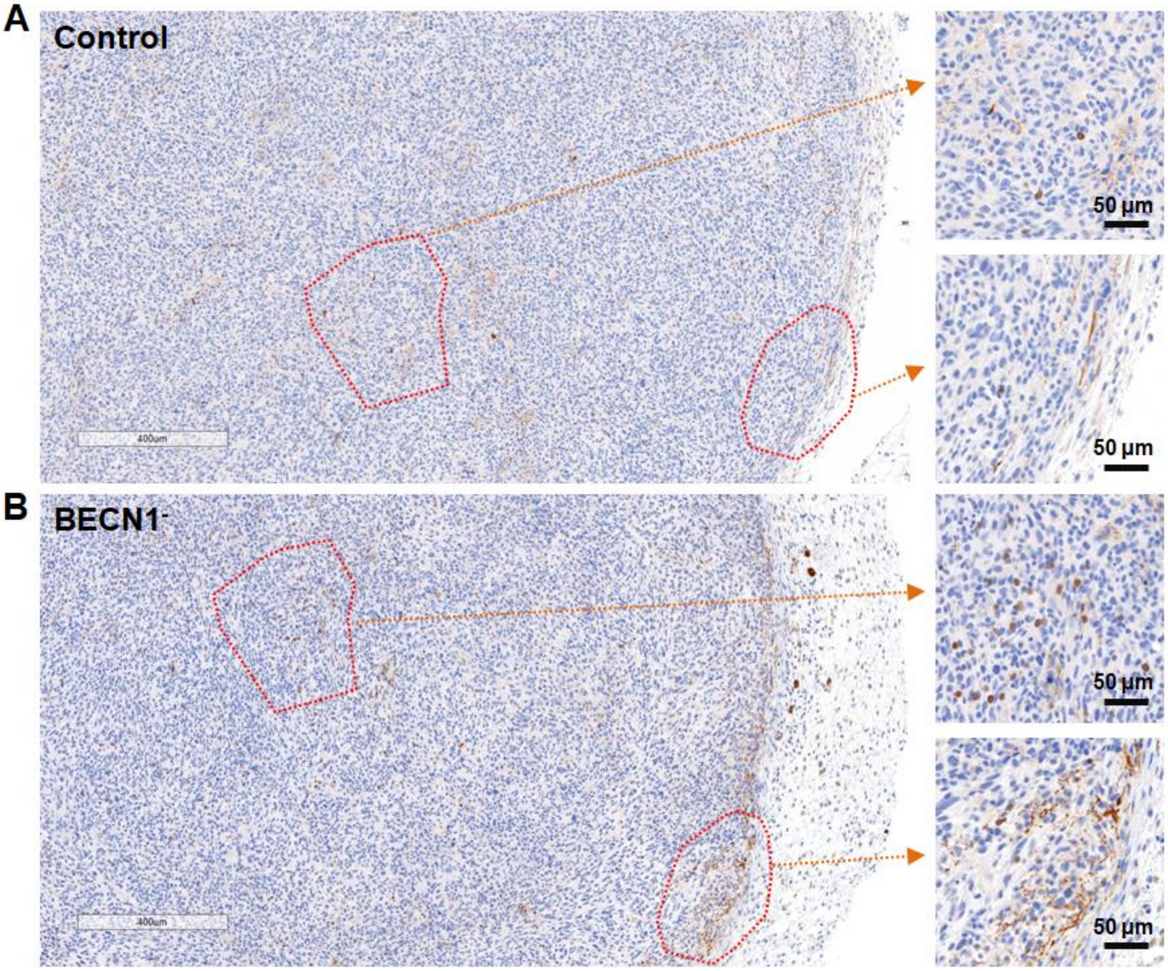


**Figure S9. Targeting autophagy in GBM through *BECN1* gene knockdown increases NK cell recruitment and homing.** (**A**) Immunohistochemical (IHC) staining of mouse ($Rag1^{-/-}$) NK cells performed on indicated GBM43 (control) subcutaneous xenograft tumor sections. (**B**) Immunohistochemical (IHC) staining of mouse ($Rag1^{-/-}$) NK cells performed on indicated *BECN1*$^{-}$ GBM43 subcutaneous xenograft tumor sections. IHC staining indicated an enhanced infiltration of NK cells into both peripheral and internal regions of the tumor tissues of *BECN1*$^{-}$ GBM43 subcutaneous xenograft tumors.

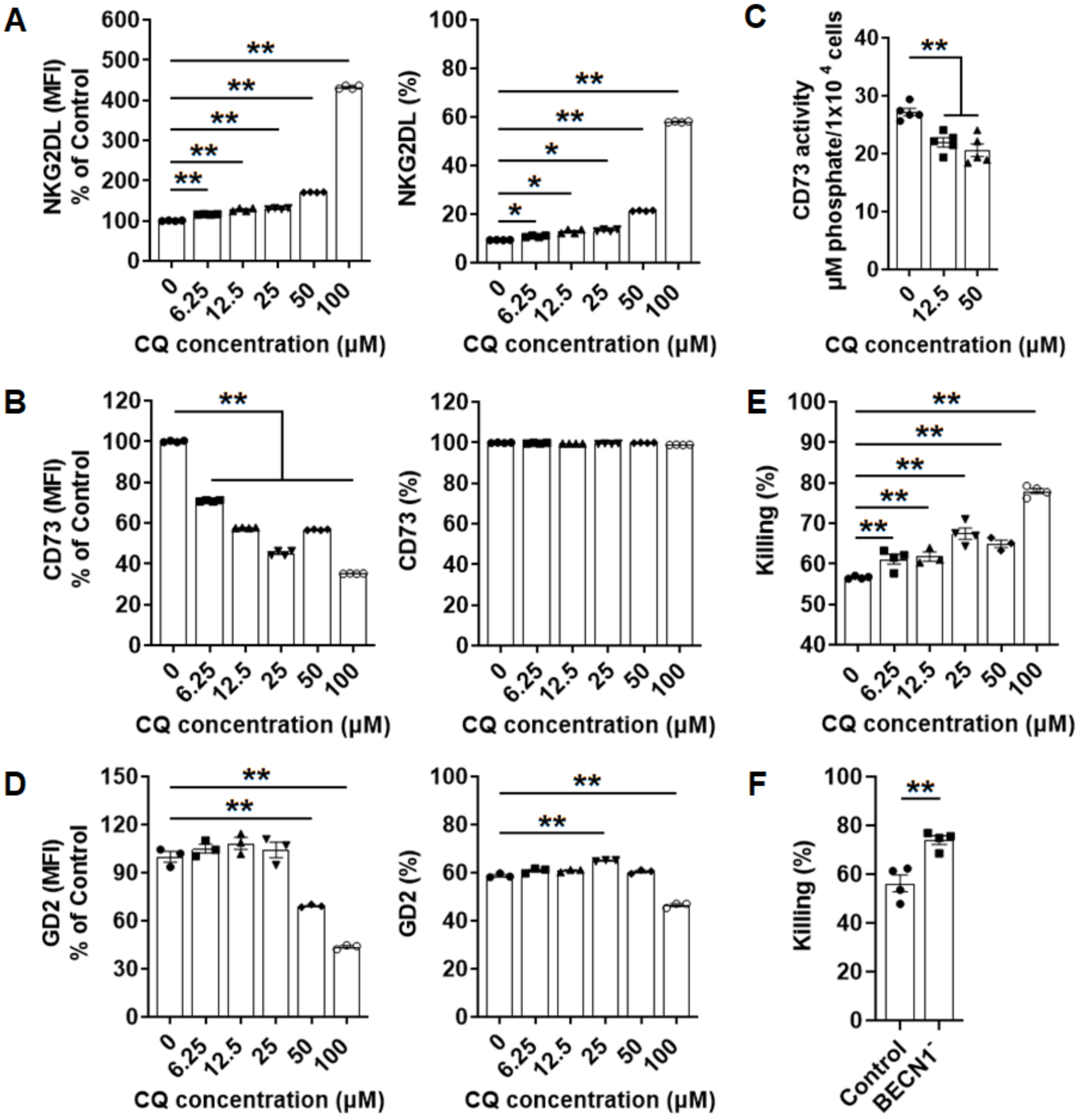


**Figure S10. Targeting autophagy sensitizes GBM cells to NK cell-mediated killing.** (**A**) NKG2DL expression (MFI and %) on GBM43 cells after 24 h treatment with various concentrations of CQ. (**B**) CD73 expression (MFI and %) on GBM43 cells after 24 h treatment with various concentrations of CQ. (**C**) CD73 activity of GBM43 cells after treatment with various concentrations of CQ for 24 h. (**D**) GD2 expression (MFI and %) on GBM43 cells after 24 h treatment with various concentrations of CQ. (**E**) Killing activity of pNK cells against GBM43 cells after treatment with different concentrations of CQ for 24 h. GBM43 cells were first treated with CQ, then collected and re-seeded into 96 wells. After overnight culture, the NK cells were added at an E/T ratio of 5 and co-incubated for 4 h. (**F**) Killing activity of pNK cells against *BECN1*$^{-}$ GBM43 cells after 4 h co-incubation at an E/T ratio of 5. Shown here are data in at least triplicates from one representative donor. Data are shown as mean ± SEM. *$p < 0.05$, **$p < 0.01$.

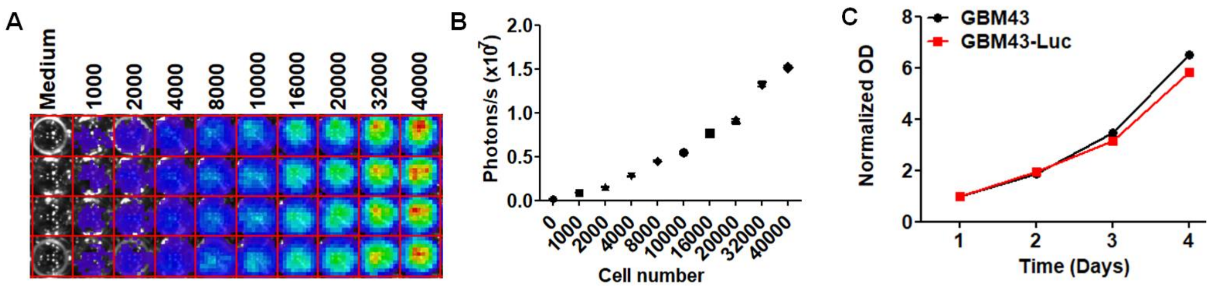


**Figure S11. Generation of luciferase-expressing GBM43 (GBM43-Luc) cells.** GBM43 cells were transfected with commercial luciferase (Luc) lentiviral particles expressing a firefly luciferase 3 gene under an inducible suCMV promoter. Puromycin-resistant cells were collected following transduction and luciferase expression was detected by bioluminescence measurement using an IVIS spectrum.(**A**) Representative bioluminescence image indicating the ability of GBM43-Luc cells to express luciferase (signal intensity as a function of cells seeded is shown). (**B**) Fluorescence intensity as a function of GBM43-Luc cell number. Total flux (photons/sec) was quantified using AURA software. (**C**) Growth curves of the GBM43-Luc and parental control GBM43 cells. Cells (1000) were grown for 4 days in regular growth medium without puromycin. Total viability of cells over time was measured using the CCK-8 assay and plotted in a logarithmic scale. Both cells showed similar growth patterns and doubling times. Data are shown as mean ± SEM.

**Video 1.** Time-lapse, over 4 hours, of killing of patient-derived GBM43 cells by engineered primary human NK (CD73.mCAR-pNK) cells. See Figure S5 for experimental details.

**Video 2.** Time-lapse, over 4 hours, of killing of patient-derived GBM43 cells by native primary human NK (pNK) cells. See Figure S5 for experimental details.
